# Multifaceted roles of a light-responsive factor LhWRKY44 in promoting anthocyanin accumulation in Asiatic hybrid lilies (*Lilium* spp.)

**DOI:** 10.1101/2023.01.09.523317

**Authors:** Mengmeng Bi, Rui Liang, Yuxiao Qu, Jiawen Wang, Yuwei Cao, Xin Liu, Guoren He, Wenliang Zhang, Yue Yang, Yuchao Tang, Panpan Yang, Leifeng Xu, Jun Ming

**Affiliations:** The Institute of Vegetables and Flowers Chinese Academy of Agricultural Sciences, Beijing, China; College of Horticulture, Shanxi Agricultural University, Taigu, Shanxi, China; College of Chemistry and Life Science, Gannan Normal University, Ganzhou, Jiangxi, China; College of Landscape architecture and Forestry, Qingdao Agricultural University, Qingdao, Shandong, China; Shanghai Key Laboratory of Plant Molecular Science, College of Life Sciences, Shanghai Normal University, Shanghai, China; College of Landscape architecture and Horticulture, Southwest Forestry University, Kunming, Yunnan, China

**Keywords:** lily, anthocyanin accumulation, LhWRKY44, MBW complex, transport regulation

## Abstract

The coloration of Asiatic hybrid lily results mostly from anthocyanin accumulation in flowers. Although anthocyanin accumulation-related genes and MBW complexes are well studied, the transcriptional regulation of WRKY transcription factors involved in anthocyanin accumulation remains poorly understood.Here, we identified a lily WRKY protein, LhWRKY44, whose expression is highly expressed downstream of LhHY5 by light and positively correlated with anthocyanin accumulation. *LhWRKY44* overexpression enhanced anthocyanin accumulation and silencing decreased anthocyanin accumulation in flowers. As a trans-acting regulator, LhWRKY44 activited the anthocyanin biosynthesis pathway-related genes *PAL* and *F3H* by binding to the their promoters. And the encoded TF also participates in anthocyanin transport and targets the intracellular anthocyanin transport protein *GST* promoters. Additionally, a novel dual activity of LhWRKY44 and LhMYBSPLATTER regulatory module, with LhWRKY44 binds to the promoter of *LhMYBSPLATTER* and interacts with LhMYBSPLATTER, strongly enhanced the interaction of LhMYBSPLATTER and LhbHLH2, indirectly enhancing *DFR, UFGT* and *GST* expression targeted by LhMYBSPLATTER. These results show a regulatory mode for light-induced anthocyanin accumulation enhancement by LhWRKY44 in lily, expanding our understanding of the complex transcriptional regulatory hierarchy modulating anthocyanin accumulation.

## Introduction

Lilies (*Lilium* spp.) are among the most common and popular ornamental plants grown worldwide with diverse floral colors. Anthocyanins, which are among the major pigments responsible for the colors of lily tepals, belongs to the flavonoid family of plant secondary metabolites and are mainly distributed in the vacuoles of plant tissues, including fruits, flowers, and leaves (Xu et al., 2017). In addition to determining the special colors of plants, these molecules also have multiple physiological functions. Such as, aid pollination, seed dispersal, and resistance to adverse environmental conditions, and medicinal functions that prevent cardiovascular disease, reduce obesity and improve glucose homeostasis in humans (Winkel-Shirley, 2001).

The anthocyanin biosynthetic pathway is the most clearly studied metabolic pathways,which involved a series of catalytic enzymes encoding the genes for synthesis. In lilies, the levels of them, e.g., *PAL*, *CHI, CHS, DFR, F3H, ANS*, and *UFGT*, directly affect anthocyanin pigmentation. LhCHS, LhF3H, LhDFR and LhANS et., have been characterized from the Asiatic hybrid lilies(Nakatsuka et al. 2003; Lai et al. 2012). In addition, there exists mechanisms for regulation of anthocyanin transport. GST, acting as a molecules carrier, could mobilizes and transported anthocyanins from the cytoplasm to the tonoplast in the process of synthetic anthocyanin being transport and storage in vacuoles (Sun et al., 2012). Xu et al. (2020) verified that the MATE family member *DTX35* is related to lily flavonoid transport. Cao et al. (2021b) revealed the anthocyanin transport mechanism mediated by *GST* in lily.

The accumulation and transport of anthocyanins is mostly regulated at the transcriptional level by the MBW complex composed of MYB, bHLH and WD40 protein. Numerous MYB proteins that act as activators or repressors have been isolated from lily, including LhMYB12 (Yamagishi et al., 2011; Yamagishi & Nakatsuka et al., 2017; Yamagishi et al., 2012; 2014a; 2014b), LhMYBSPLATTER-Latvia (Yamagishi et al., 2014a;b), LhMYB6 (Yamagishi et al., 2010), LhMYB18 (Yamagishi, 2018), LhR3-MYB (Sakai et al., 2019), LhMYB19Long (Yamagishi, 2020a; 2020c) and LrMYB15 (Yamagishi, 2016). Addition, LhbHLH1, LhbHLH2, LhWD40a and LhWDR were characterized and involved in anthocyanins accumulation in lily(Nakatsuka et al., 2008; Yamagishi et al., 2010;Suzuki et al., 2016; Dou et al., 2020). Besides, TFs such as HY5, SPL, NAC, AP2/ERF, HD-Zip, BBX, and WRKY were found regulating anthocyanin accumulation in plants recently (Li et al., 2020; Ma et al., 2021). They regulate anthocyanin accumulation by directly or indirectly functions with structural genes or MBW complexes. Pp4ERF24 and Pp12ERF96 regulate anthocyanin accumulation in pear fruit by physically interacting with PpMYB114 (Ni et al., 2019). The FaRAV1 binds to the *FaCHS, FaF3H, FaDFR* and *FaGT1* promoters to promote anthocyanin biosynthesis in strawberry fruit (Zhang et al., 2020).

WRKY superfamily members are characterized by WRKY domains and zinc-finger motif. They can recognize and bind to a specific W-box sequence, (C/T) TGAC (T/C), to play regulatory roles (Zentgraf et al., 2010). It has been extensively shown that members of the WRKY family participate in anthocyanin accumulation-associated processes in other species. In particular, *Arabidopsis TTG2* encodes a WRKY44 protein, which affects flavonoid accumulation (Gonzalez et al., 2016; Verweij et al., 2016). In apple, *MdWRKY11* promoted anthocyanin accumulation in the callus by enhancing transcriptional activation of *MdMYB10, MdMYB11* and *MdHY5* (Liu et al., 2019). MdWRKY40, a wounding-responsive protein, is involved in apple peel anthocyanin biosynthesis by interaction with MdMYB1, and activate *MdF3H*, *MdUF3GT*, *MdCHS* and *MdCHI* expression (An et al., 2019). In pear, PyWRKY26 interacts with PybHLH3 and binds to the *PyMYB114* promoter to promote anthocyanin accumulation (Li et al., 2020). Although abundant evidences showed WRKY TFs participare in anthocyanin accumulation in plants, no studies on the regulation by the lily WRKY family have been reported, and the regulatory mechanisms are currently unknown.

In previous study, we screened all annotated WRKY genes’expression during flowers development based on transcriptome data, and identified a lily WRKY gene(c117585—g2), LhWRKY44 closely related to anthocyanin accumulation in lily Tango series cultivar ‘Tiny Padhye’. Here, we identified a light-responsive factor LhWRKY44 positively regulates anthocyanin biosynthesis via the transcriptional regulation of *PAL*,*F3H*, and a intracellular anthocyanin transport protein *GST*. Additionally, we also demonstrated a novel WRKY-MBW module different from those of PH3 and TTG2 where LhWRKY44 binds to the *LhMYBSPLATTER* promoter and interacts with LhMYBSPLATTER. The interaction between LhMYBSPLATTER and LhbHLH2 is strongly enhanced and indirectly enhances the expression of *DFR*, *UFGT* and *GST*, which are targeted by LhMYBSPLATTER, thus influencing anthocyanin accumulation. Collectively, Our findings have further clarified the effects of LhWRKY44-mediated transcriptional regulation on anthocyanin in lily.

## Results

### Isolation and characterization of the LhWRKY44

The full-length 1972-bp LhWRKY44 cDNA was isolated from ‘Tiny Padhye’ by RACE. The segments comprised 1386 bp open reading frames encoded a 461-residue polypeptide. Sequence analysis showed that LhWRKY44 protein possessed two conserved WRKYGQK domains toward the N-terminus and a zinc finger ligand. (Supplemental Figure S1). Cluster analysis indicated that LhWRKY44 was most closely to a branch containing AtWRKY44 (TTG2), hence its designation as LhWRKY44 (Supplemental Figure S2). LhWRKY44 belongs to group I of the WRKY superfamily. Phylogenetic analysis of WRKY proteins confirmed to be associated with anthocyanin accumulation from multiple species showed that LhWRKY44 shares a close evolutionary relationship with PhPH3 (WRKY44) of *petunia* (Figure 1A).

**Figure 1.**
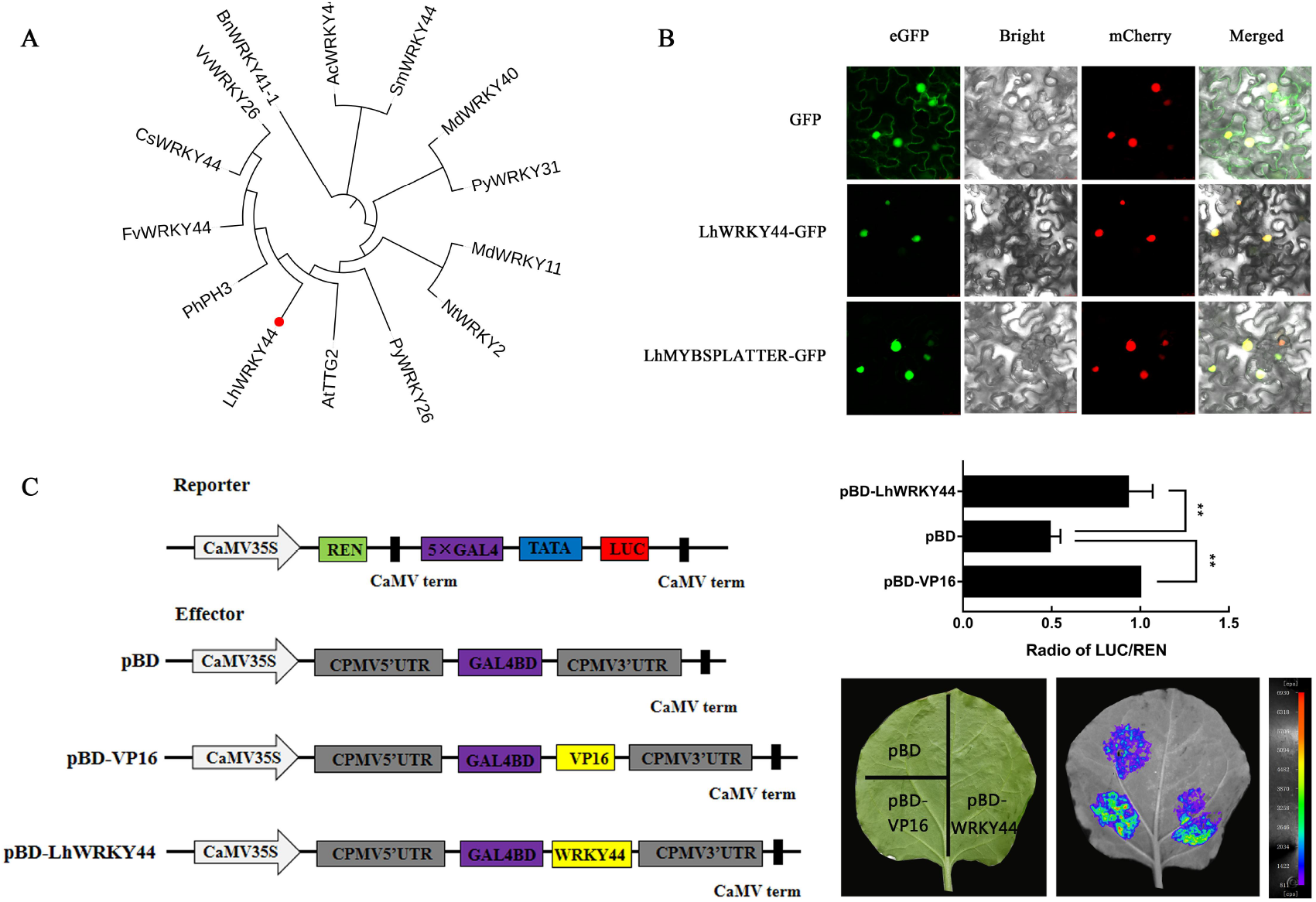
Characterization of the LhWRKY44 transcription factor. A,Phylogenetic analysis of LhWRKY44 and anthocyanin-related WRKY proteins from other plant species. The amino acid sequences of SmWRKY44 (AKA27910.1), PhPH3 (AMR43368.1), AtTTG2 (NP_181263.2), CsWRKY44 (AYA73391), VvWRKY26 (AQM37647), BnWRKY41-1 (XP_013686534.1), PyWRKY26 (Pbr013092.1), PyWRKY31 (Pbr000122.1), FvWRKY44 (XM_004302784), NtWRKY2 (AB_063576), MdWRKY11 (MDP0000128463), AcWRKY44 (ACC16887.1), and MdWRKY40 (XP_008342807.1) were retrieved from the GenBank database. B,Subcellular localization of the LhWRKY44-GFP and LhMYBSPLATTER-GFPs in *N. benthamiana* leaf epidermal cells with the nuclear marker mCherry. eGFP, GFP signal; mCherry, nuclear marker; Bright, white light; Merged, combined GFP and mCherry signals. Scale bars, 25 μm. C,Transcriptional activation activity of LhWRKY44 in *N. benthamian*a leaves. Schematic diagram of vector left).pBD-LhWRKY44, the recombinant plasmid containing LhWRKY44; pBD, the negative control; pBD-VP16, the positive control. live imaging and quantitative analysis (right). Mean values ± SDs are shown at least three biological replicates. Asterisks represent statistically significant differences (*, P < 0.05; **, P < 0.01; ***, P < 0.001,T test and ANOVA).

To confirm LhWRKY44 putative function as a TF, we generated a LhWRKY44-GFP recombinant protein and conducted subcellular localization assay. The fluorescence signal matched the GFP signal (Figure 1B), indicating that LhWRKY44 localizes to the nucleus. A further transcriptional activity in *N. benthamiana* leaf cells revealed that the relative luciferase in pBD-LhWRKY44 exhibited much higher activity than the negative control pBD (Figure 2C). Thus, LhWRKY44 work as a nucleus-localized transcriptional activator.

**Figure 2.**
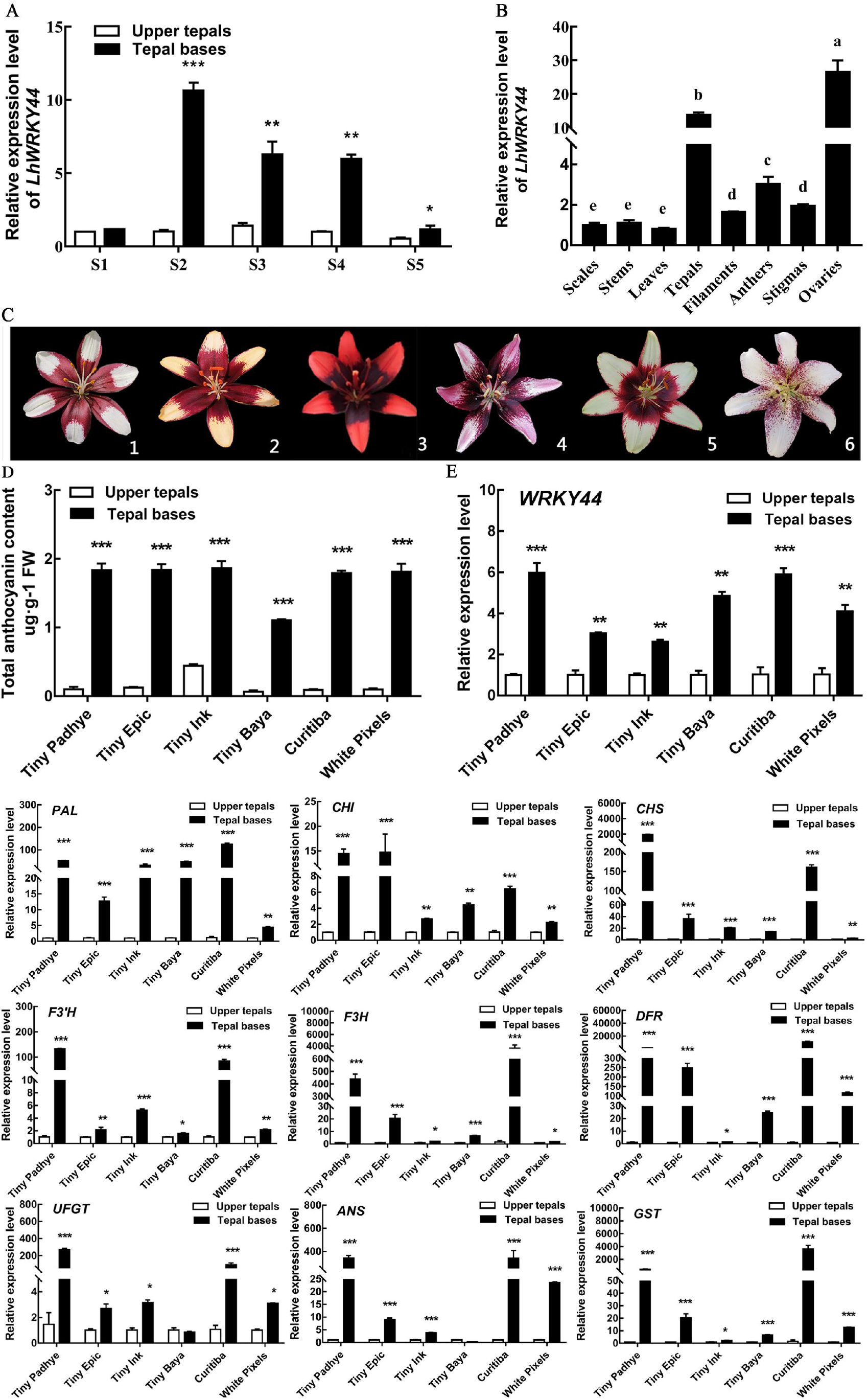
The transcription level of *LhWRKY44* is related to anthocyanin accumulation in lily. A,Expression of *LhWRKY44* in different developmental stages of ‘Tiny Padhye’ tepals.B,Expression of *LhWRKY44* in different tissues of ‘Tiny Padhye’. C,Tango series lily cultivars. 1: ‘Tiny Padhye’; 2: ‘Tiny Epic’; 3: ‘Tiny Ink’; 4: ‘Tiny Baya’; 5: ‘Curitiba’; 6: ‘White Pixels’. D,Determination of the anthocyanin content. E, Relative transcript levels of *LhWRKY44*, anthocyanin biosynthetic genes and regulatory genes. Flowers (stage 3) were subjected to RNA extraction and transcriptome analysis.The lily *actin* gene was used as the control. Mean values ± SDs are shown at least three biological replicates. Asterisks represent statistically significant differences (*, P < 0.05; **, P < 0.01; ***, P < 0.001,T test and ANOVA).

### Expression of *LhWRKY44* is closely associated with anthocyanin accumulation

RT-qPCR analysis result at different stages of development showed *LhWRKY44* was always higher in basal tepals than in upper tepals, in accordance with the anthocyanin content (Figure 2A; Supplemental Figure S3). And the expression levels of *LhWRKY44* were higher in the flowers and ovaries tissues than in anthers, filaments, stigmas, stems, scales and leaves in tissue specific expression pattern analysis (Figure 2B; Supplemental Figure S3).The relatively high expression of *LhWRKY44* indicated that LhWRKY44 is spatially and temporally connected with lily anthocyanin accumulation.

To further confirm the reliability of the expression of LhWRKY44 is closely associated with anthocyanin accumulation, five other Tango series cultivars of Asiatic hybrid lilies considered to be associated with the color type caused by anthocyanin accumulation (Yamagishi, 2013b) were chosen to determine the expression levels of *LhWRKY44* in basal (high anthocyanin level) and upper (low anthocyanin level) tepals at the same developmental stages by RT-qPCR (Figure 2C). The results further demonstrated that the *LhWRKY44* level increased with deepening of the tepal color and was higher than that in the upper areas, consistent with the anthocyanin levels and the levels of *LhCHI, LhCHS, LhF3H, LhF3’H, LhDFR, LhUFGT, LhANR*, and *LhANS* (Figure 2D and E). These data reveal that the expression of *LhWRKY44* showed a significant positive correlation with the anthocyanin content and may be related to anthocyanin accumulation.

### Overexpression of *LhWRKY44* leads to a stronger red coloration and silencing leads to a weaken red coloration in lily tepals

To investigate the functions of LhWRKY44 in lily tepal anthocyanin accumlation, transient transformation experiments were carried out. Agrobacterium cultures harboring 35S:GUS (empty vector) and 35S:LhWRKY44 were used to infiltrate the outer tepals of lily cultivar ‘Robina’ buds. Indeed, we found that the control group and overexpression group definitively showed a few changes in flower color, as expected (Figure 3A-C; Supplemental Figure S4A and B). However, when tepals from the same stage of the lily buds that showed color were infected with the recombinant TRV construct, the color of those infected with the pTRV2-LhWRKY44 vector was clearly lighter than that of the control (Figure 3A-C, Supplemental Figure S4C and D).

**Figure 3.**
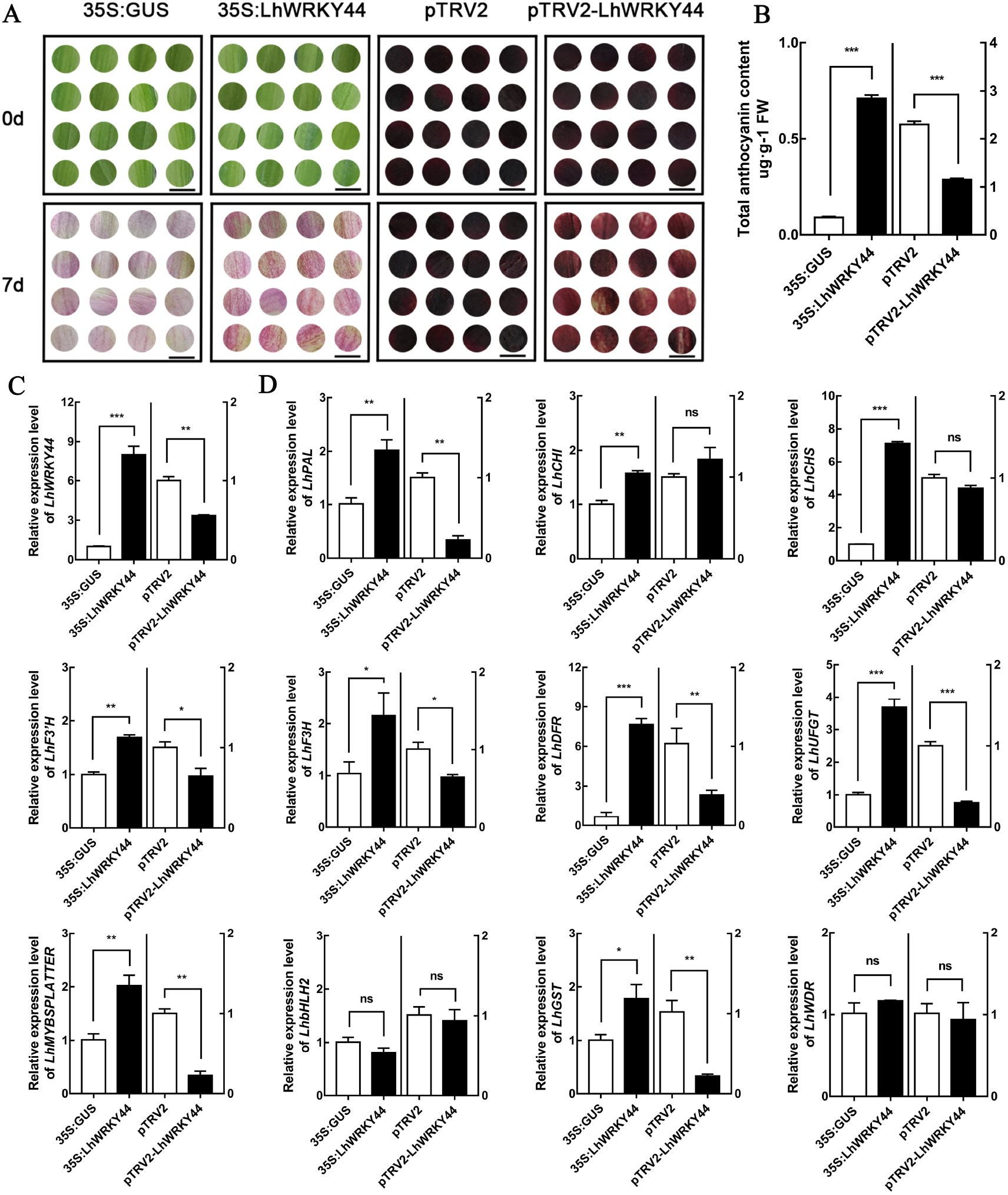
Transient overexpression and silencing of *LhWRKY44* in lily tepals. A,Phenotypic images of the transient overexpression and silencing of *LhWRKY44* in the infiltrated sites (approximately 1 cm of tepals). Phenotypic images taken 7 d after infiltration. Bar=1 cm. B,Anthocyanin content after overexpression and silencing of *LhWRKY44*. C,Expression levels of *LhWRKY44* in lily tepals with transient expression and silencing. D,Expression of anthocyanin accumulation-related genes in lily tepals transiently expressing *LhWRKY44* or with the genes silenced. Mean values ± SDs are shown at least three biological replicates. Asterisks represent statistically significant differences (*, P < 0.05; **, P < 0.01; ***, P < 0.001,T test and ANOVA).

Further analysis showed that the expression of *LhWRKY44* had different effects on structural genes involved in anthocyanin accumulation and anthocyanin regulation-related genes. For instance, *PAL* transcript levels were markedly higher in the group transiently overexpressing *LhWRKY44* than in the control but much lower in the pTRV2-LhWRKY44 treatment group than in the pTRV2 empty group. *F3’H*, *F3H, DFR* and *UFGT* showed a similar expression pattern as that of *PAL* (Figure 3C). Beside, the expression levels of *LhMYBSPLATTER* clearly decreased in pTRV2-LhWRKY44 compared with the control, while a nearly 6-fold increase was recorded in the overexpression group compared with the control. *LhMYB6* and *LhGST* were moderately increased,but no difference on *LhbHLH2 and LhWDR* (Figure 3D; Supplemental Figure S4E). In summary, LhWRKY44 promotes anthocyanin accumulation in lily tepals, and may regulates anthocyanin accumulation-related genes.

### LhWRKY44 activates the expression of anthocyanin pathway genes by directly targeting their promoters

Considering that the transcript levels of anthocyanin pathway genes were significantly upregulated after overexpressed, these genes may be direct targets of LhWRKY44. To prove this conjecture, we first analyzed the FRKM values of the tepals in different periods. Five anthocyanin pathway genes *LhPAL, LhCHS, LhF3H, LhDFR* and *LhUFGT* were consistent with the expression levels of *LhWRKY44*. Thus, we isolated the promoters and construct the vector for dual-luciferase assay(Figure 4A; Supplemental Figure S5). A transiently transformed *N. benthamiana* leaves was carried out to investigate the effects of LhWRKY44 on promoters activity. As shown as Figure 4A and B, LhWRKY44 activated the transcription of these genes to different extents, with the highest expression observed for the *LhPAL* and *LhF3H* promoter, followed by *proLhDFR, proLhCHS* and *proLhUFGT*.

**Figure 4.**
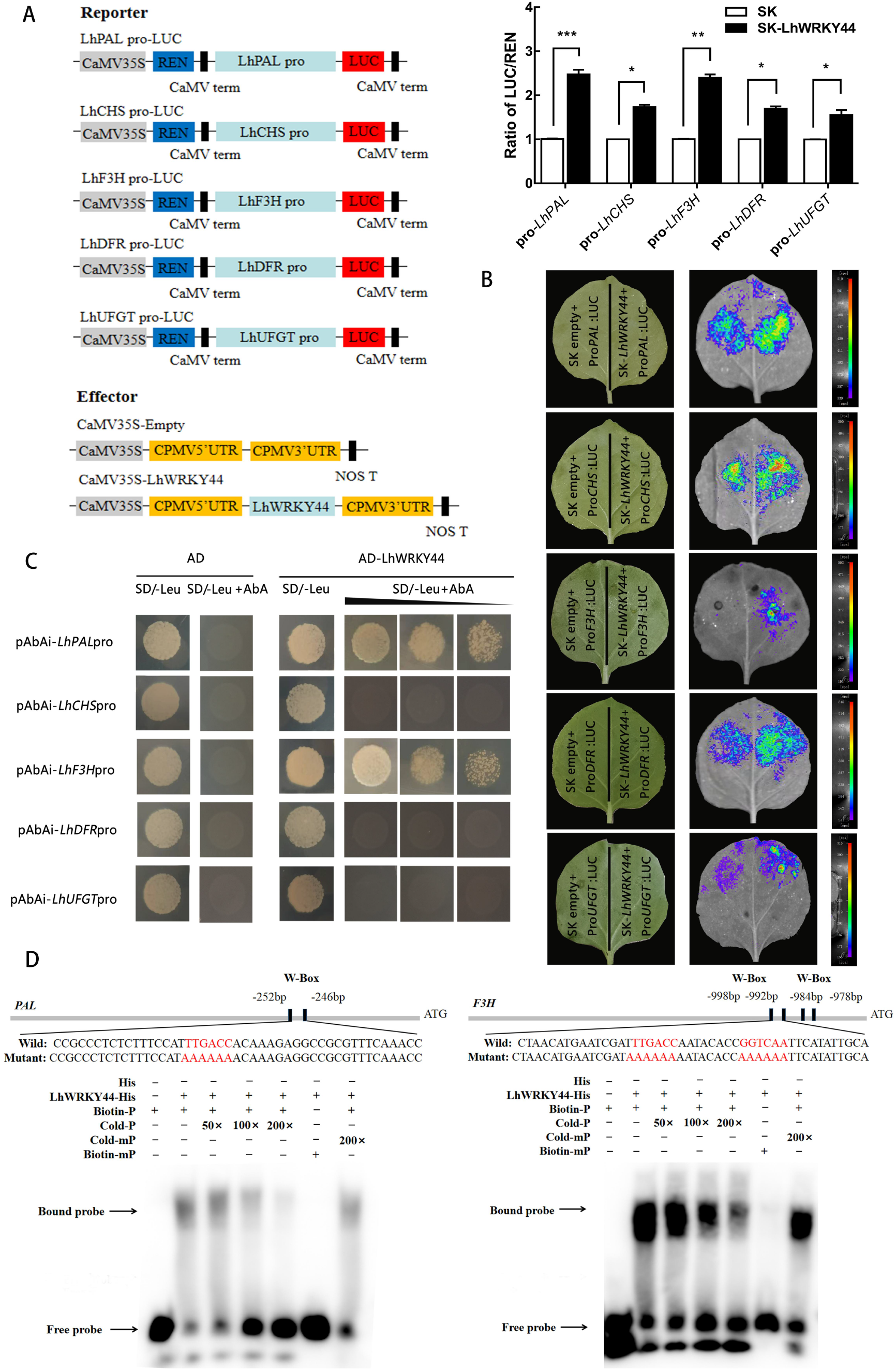
LhWRKY44 activates anthocyanin pathway genes’ expression and binds to *LhPAL* and *LhF3H* promoters. A,Dual-luciferase analysis of the effect of LhWRKY44 on the promoters of anthocyanin biosynthesis genes. Diagrams of the reporter and effector vectors of the dual-luciferase transient expression assay (left) and the LUC/REN ratio (right).SK refers to empty vector and is set as 1. B,Schematic diagram of the injected *N. benthamiana* leaves and living images by CDD. Mean values ± SDs are shown at least three biological replicates. Asterisks represent statistically significant differences (*, P < 0.05; **, P < 0.01; ***, P < 0.001,T test and ANOVA).C,Yeast one-hybrid analysis of the interaction of LhWRKY44 and anthocyanin biosynthetic gene promoters. D,LhWRKY44 binds to the W-box element of LhPAL and LhF3H in EMSAs. The probe sequences are shown, The term 250 × indicates the usage of excess cold competition probe, and “+” and *“-”* indicate its presence and absence, respectively. Arrows indicate the positions of protein-DNA complexes or free probes.

To confirm the direct binding of LhWRKY44 to the target promoters, Y1H assays was performed. The basal activity of promoters were detected in yeast in the presence of aureobasidin A (AbA). The yeast cells cotransformed with LhWRKY44 and the negative control all survived on SD/-Leu. However, only the yeast cells cotransformed with LhWRKY44 and *PAL* promoter or *F3H* promoter could survive on SD/-Leu/AbA (Figure 4C). Thus, LhWRKY44 protein could interact with the *PAL* and *F3H* promoters in the yeast system. WRKY proteins target promotersgenes by recognizing a specific DNA sequence, the W-box ((T) GAC (C/T)), (An et al., 2019). Analysis of the promoters showed the presence of at least one W-box motif in *PAL* and *F3H* promoters (Supplemental Figure S5). Then, EMSAs were conducted and specifically binding of LhWRKY44 to W-box was characterized in *PAL* and *F3H* promoters. As shown in Figure, 4D. A mobility shift was produced when LhWRKY44-His proteins were incubated with the W-box probe, and impaired with the adding of unlabeled competitor DNA probe but recovered with the addition of mutated unlabeled DNA probe.These data support the notion that LhWRKY44 acts as a transcriptional activator of anthocyanin accumulation pathway genes by directly binding to their promoters.

### LhWRKY44 participates in anthocyanin transport by binding *LhGST* promoters

Anthocyanin is catalyzed by the multi-enzymes in the cytoplasm, and then transported important to the vacuole. It has been proved that members of the glytide transferase (GST) family played key roles in the transportation of anthocyanins. To date, LhGST have been implicated in anthocyanin transport in lily (Cao et al., 2021).

Considering that the transcript levels of *LhGST* were also increased in *LhWRKY44* overexpressing tepals and down-regulated after silencing LhWRKY44 (Figure 3D). Besides, one W-box motif was found within the region upstream of the start codon after searching the promoter of *LhGST* (Supplemental Figure S5). So we then raise the question of whether *LhGST* was also the direct targets of LhWRKY44. Dual-luciferase assay showed a activation of *LhGST* promoter (Figure 5A). Based on autoactivation, three truncated regions of the *LhGST* promoter fragment P1, P2 and P3, were used as baits and assayed with the LhWRKY44 protein. LhWRKY44 bound the P1 containing W-box motif and did not bind the P2 or P3 without W-box(Figure 5B). Results of electrophoresis mobility shift assay (EMSA) further supported aboved result (Figure 5C). Consequently, these results provided evidences for LhWRKY44 could participates in anthocyanin transport in lily by binding *LhGST* promoters.

**Figure 5.**
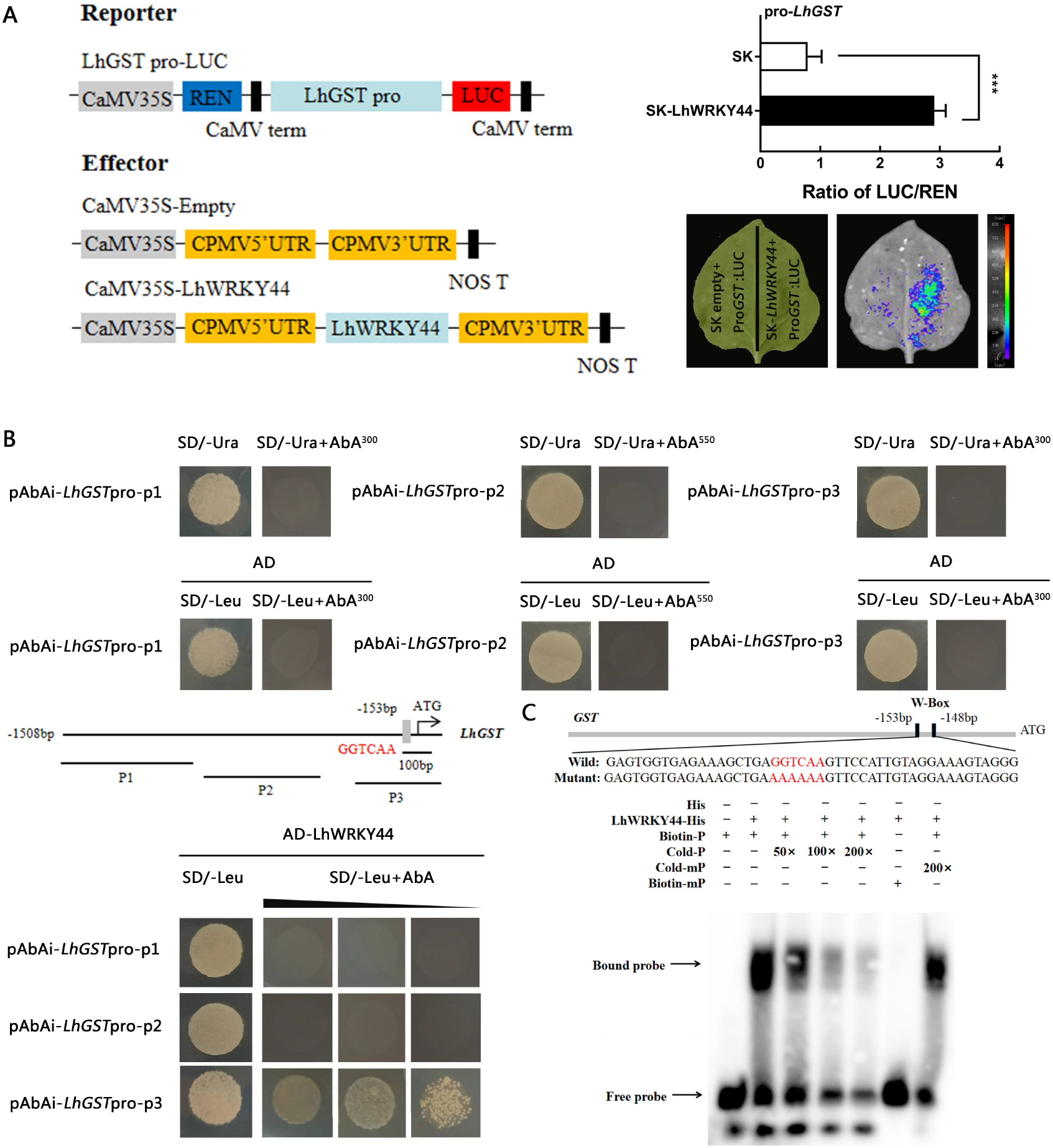
Direct binding of LhWRKY44 to the promoters of *LhGST*. A, Dual-luciferase assay showing activation of LhWRKY44 to *ProGST*. Mean values ± SDs are shown at least three biological replicates. Asterisks represent T test significance (*P < 0.05,**P < 0.01 and ***, P < 0.001). B,Yeast one-hybrid analysis of the interaction of LhWRKY44 and *LhGST* promoters. C, LhWRKY44 binds to the W-box element of *LhGST* promoter in EMSAs.

### LhWRKY44 physically interacts with LhMYBSPLATTER

Many other TFs may interact with MBW complexes or function upstream to regulate their functions (Xu et al., 2014). To clarify whether LhWRKY44 regulates anthocyanin biosynthesis via MBW complex, we identified if LhMYBSPLATTER, LhbHLH2 and LhWDR as LhWRKY44 potential interaction partners, individually. Directed BiFC assays validated that LhWRKY44 interacted with LhMYBSPLATTER, but not with LhbHLH2 and LhWDR in vivo (Figure 6A). The fluorescence signal generated upon LhWRKY44 interacting with LhMYBSPLATTER was observed in the nucleus when tobacco leaves were coinfiltrated with LhWRKY44-YFPC and LhMYBSPLATTER-YFPN. There is a lack of fluorescence upon coinfiltration with LhWRKY44-YFPC and YFPN or YFPC and LhMYBSPLATTER-YFPN. So were the combinations of LhWRKY44 with LhbHLH2 or LhWDR. Subsequently, the interaction relationship was proven by a Y2H assay. The Y2H assay results further supported aboved result (Figure 6B). FLC imaging assay also showed that LhWRKY44 could interact with LhMYBSPLATTER (Figure 6C). Notably, of these, LhWRKY44 was also proved could activites the expression of *LhMYBSPLATTER* and binds to the W-box motif on the *LhMYBSPLATTER* promoter (Supplemental Figure S8). These results further revealed a novel dual role of WRKY-MBW between LhWRKY44 and LhMYBSPLATTER controls anthocyanin accumulation in lily.

**Figure 6.**
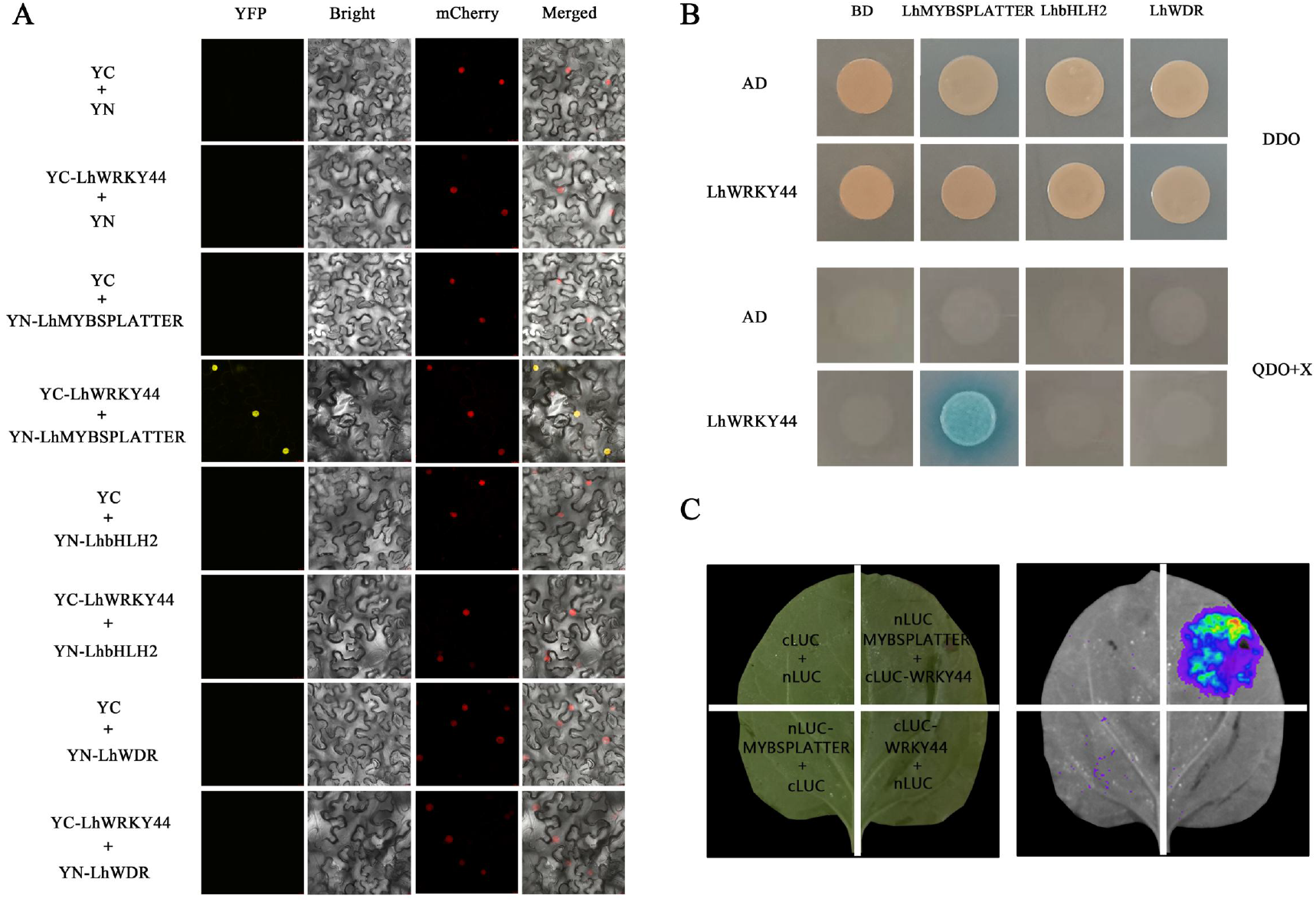
LhWRKY44 physically interacts with LhMYBSPLATTER. A,BiFC assays of LhWRKY44 and MBW complex in *N. benthamiana* leaf epidermal cells B,Yeast two-hybrid assay showed LhWRKY44 interacts with LhMYBSPLATTER among MBW complex.C, FLC assay. LUC signals were imaged using a CCD camera.

### LhWRKY44 enhances the interaction between LhMYBSPLATTER and LhbHLH2

To examine the effect of the interactions between LhWRKY44 and LhMYBSPLATTER on the MBW complex, we conducted an FLC imaging assay. LhMYBSPLATTER and LhbHLH2 fused to the N and C termini of luciferase, respectively. And LhWRKY44 was inserted into the pGreenII 0029 62-SK vector. Tobacco leaves were coinfiltrated with different combinations of the recombination plasmids. As shown as Figure 8A, the luminescence was observed in regions containing LhMYBSPLATTER-cluc and LhbHLH2-nluc, but not in regions with LhMYBSPLATTER-cluc and empty-nluc. When co-infiltration of LhWRKY44-SK in a 1:1:1 ratio to LhMYBSPLATTER-cluc and LhbHLH2-nluc enhanced the luminescence, and co-infiltration of LhWRKY44-SK in a 5:1:1 ratio to LhMYBSPLATTER-cluc and LhbHLH2-nluc strongened the luminescence. Results suggested that LhWRKY44 enhanced the interaction between LhMYBSPLATTER and LhbHLH2, promoted with the formation of MBW complex.

**Figure 7.**
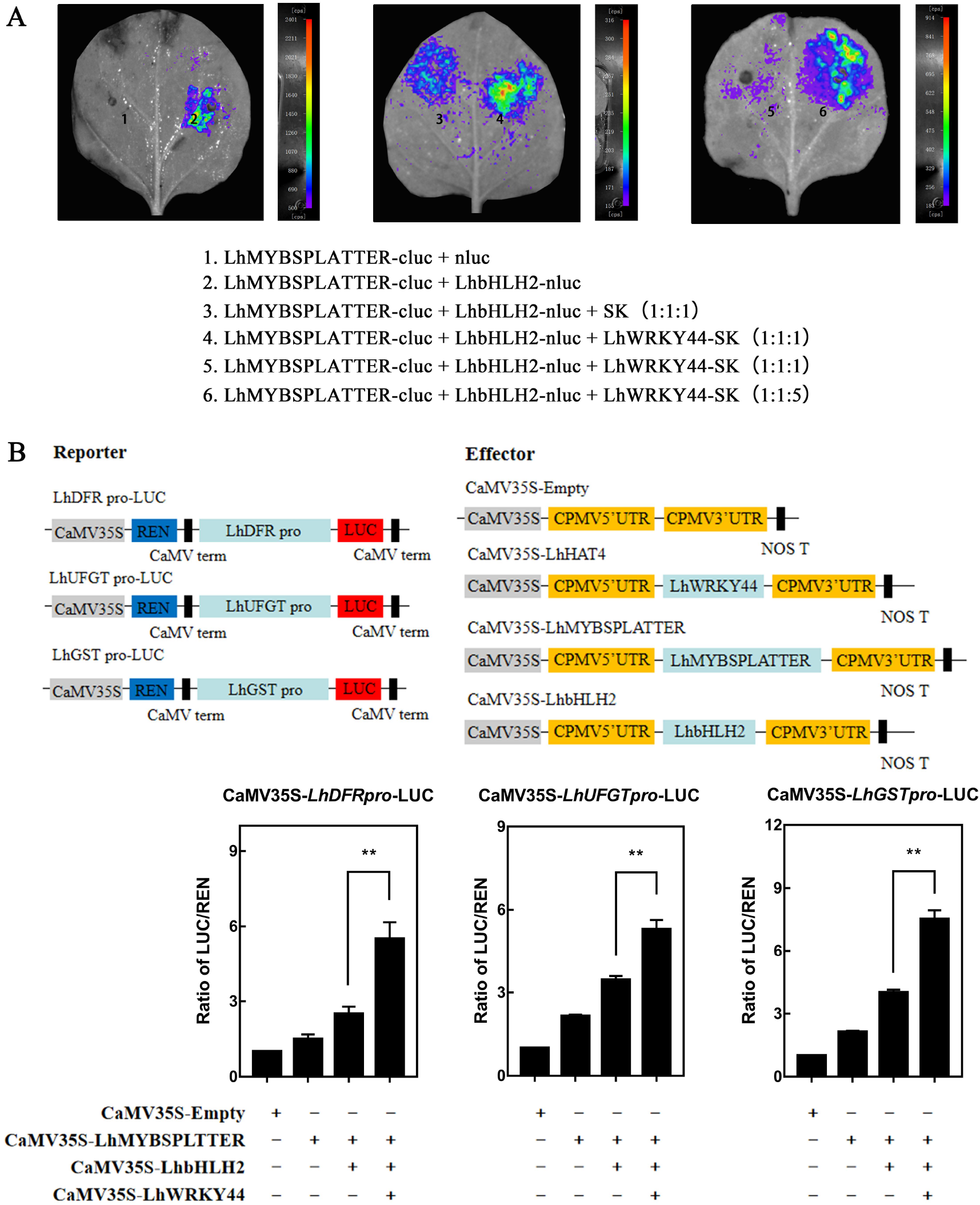
Effects of LhWRKY44 on the interaction between LhMYBSPLATTER and LhbHLH2. A, FLC imaging assays. B,Schematic diagram of tobacco leaf injection. A dual-luciferase assay verified that cotransformation with different combinations of LhWRKY44, LhMYBSPLATTER and LhbHLH2 affected *DFR*, *UFGT* and *GST* promoter activity. Mean values ± SDs are shown at least three biological replicates. Asterisks represent T test significance (*P < 0.05,**P < 0.01 and ***, P < 0.001).

**Figure 8.**
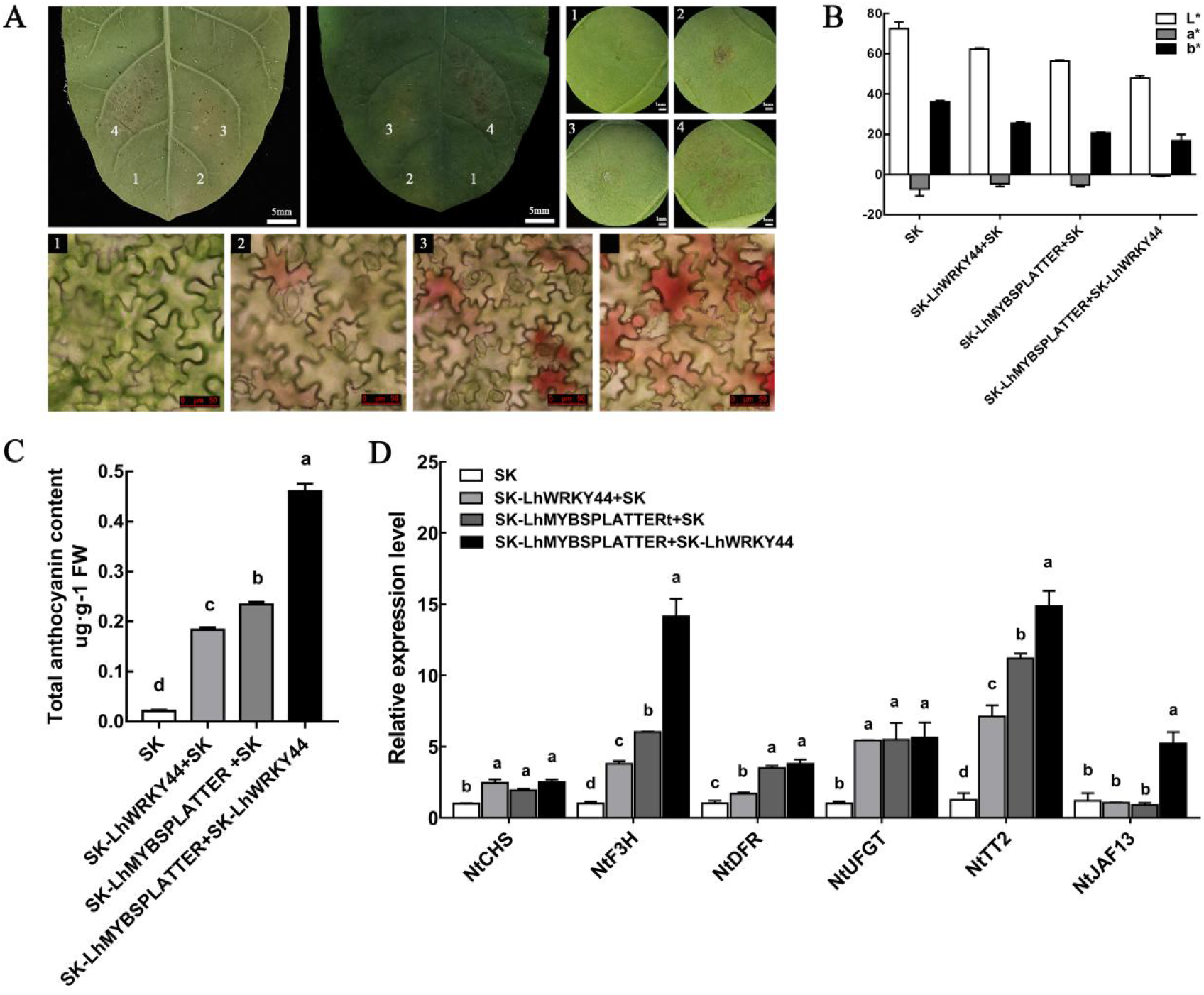
Coexpression of *LhWRKY44* and *LhMYBSPLATTER* in *N. tabacum* leaves.A,Color development photographs after infiltration in *N. tabacum*. B, Measurement of the a*/b* ratio in colored regions by a Minolta Chroma Meter. C, Anthocyanin content of tobacco leaves. D,Expression of anthocyanin biosynthetic genes and regulatory genes in *N. tabacum*.Mean values ± SDs are shown at least three biological replicates. Different letters indicate significantly different values (P<0.01) using ANOVA followed by a Tukey’s correction. Different letters are significantly different (P<0.01, ANOVA, Tukey’s correction).

Previously, we screened LhMYBSPLATTER functions as a coregulator of anthocyanin accumulation with LhbHLH2 by directly targeting the promoters of *DFR*, *UFGT* and *GST*, which are anthocyanin regulation-related genes(Yamagishi et al., 2010; Cao et al., 2021b; unpublished). Therefore, the LhWRKY44 - LhMYBSPLATTER complex might affect the interactions of LhMYBSPLATTER to the *DFR*, *UFGT* and *GST*. We tested this hypothesis by a dual-luciferase assay. As shown in Figure 8B. LhWRKY44 was observed to upregulate *DFR, GST* and *UFGT* expression at different levels in the presence of LhMYBSPLATTER and LhbHLH2. The LUC/REN ratio was significantly higher than that observed when only LhMYBSPLATTER or LhbHLH2 was expressed, which suggests that the LhWRKY44-LhMYBSPLATTER interaction promoted the ability of LhMYBSPLATTER to activate its target genes *DFR*, *UFGT* and *GST*.

A transient transformation assay involving *N. tabacum* leaves as described in Xiang (2019) was also conducted. As shown in Figure 9A, no pigmentation was observed at the infiltration sites of leaves with SK (empty vector) alone. However, anthocyanin accumulation could be weakly triggered by LhWRKY44 or LhMYBSPLATTER alone at 8 d. The addition of LhWRKY44 in LhMYBSPLATTER lead to the formation of a deeper color at an earlier time point 3 d after infiltration (Figure 8A-C). The structural genes and regulatory genes (*NtCHS, NtDFR*, *NtUFGT*, and so on) associated with anthocyanin accumulation were significantly upregulated in the leaves upon cotransformation of the two TFs (Figure 8D). Overall, these results clearly demonstrated that LhWRKY44 promoted the forming and interaction of MBW complex. The activited formation of the MBW complex increases anthocyanin accumulation and the transcriptional activation in cells.

**Figure 9.**
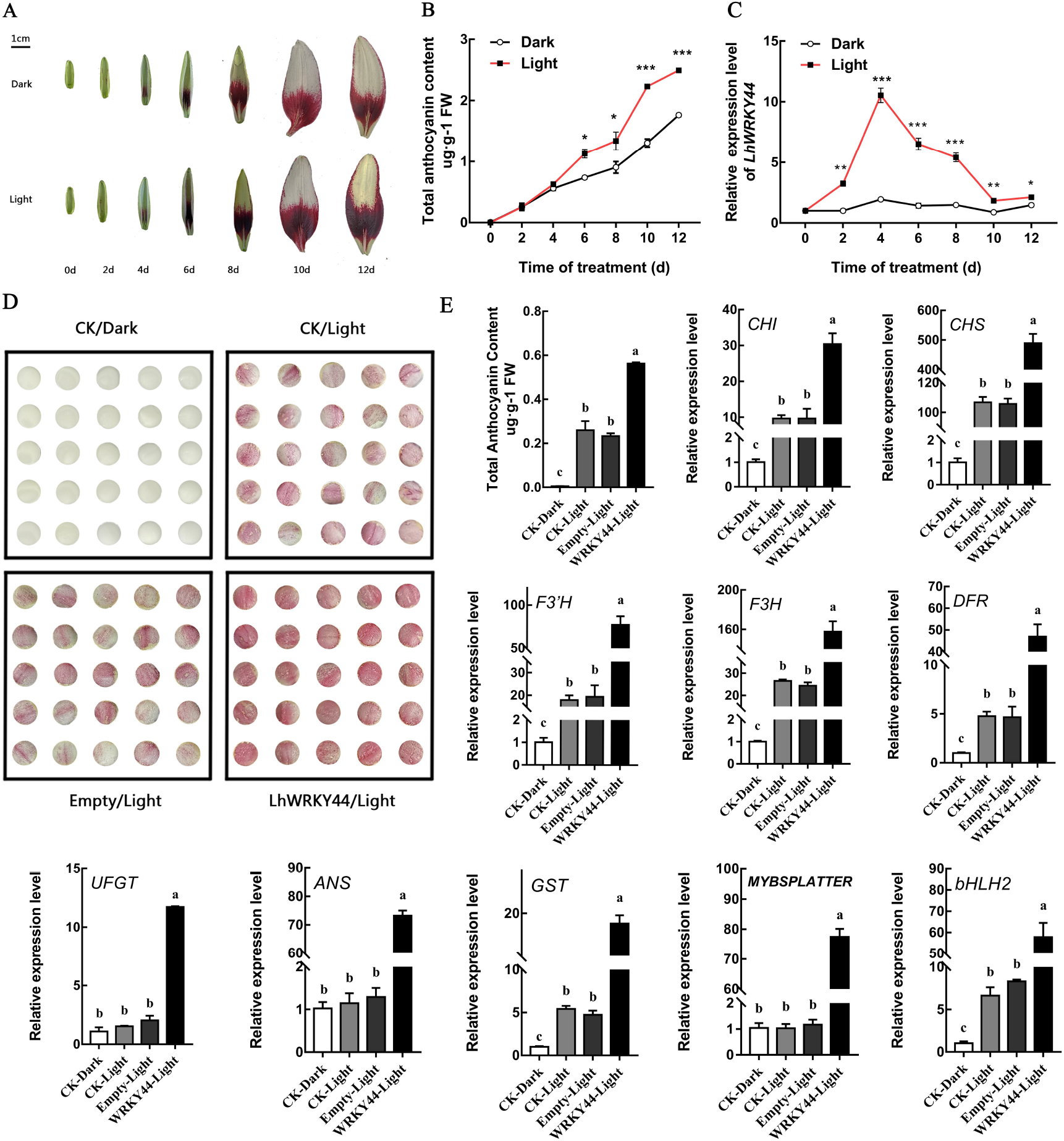
LhWRKY44 is related to light-induced anthocyanin accumulation in lily. A, Effects of light on flower color. Flower buds ‘Tiny Padhye’ at different stages of under dark and light treatment for different lengths of time. Scale bar, 1 cm. B, Anthocyanin content analysis treated with high light for different durations.C,Relative expression of *LhWRKY44*.D,Scale color development photographs after infiltration under dark and light treatments. CK: control, water treatment; Empty: empty vector treatment; LhWRKY44: transient overexpression of *LhWRKY44* treatment.E, Anthocyanin content and qPCR analysis. All samples were collected at 4 d after treatment. Mean values ± SDs are shown at least three biological replicates. Asterisks represent T test significance (*P < 0.05,**P < 0.01 and ***, P < 0.001). Different letters indicate significantly different values (P<0.01) using ANOVA followed by a Tukey’s correction.

### *LhWRKY44* is involved in light-induced anthocyanin biosynthesis in lily

Considering that there are a large number of light-responsive elements in *LhWRKY44* promote,we tested whether the expression of *LhWRKY44* could be induced by light. Lily plants were subjected to high-light (60000 lux) and weak-light (dark) treatment. As shown as Figure 9A, the anthocyanin accumulation significantly differed between the light-treated and control groups only after 6 d of the treatment period when flower buds were ready to open, and increasing light treatment times were associated with increasing intensity of the red coloration of lily tepals. The transcript levels of LhWRKY44 were highly upregulated (within 12 d) after exposure to light and peaked 4 d after treatment (Figure 9B). In contrast, the expression in the control hardly changed over time. Anthocyanin accumulation-related genes were more highly expressed in the light-treated samples than in the control to various degrees (Supplemental Figure S9), except *LhMYBSPLATTER* at 10 d and *LhWDR* at 6 d.

Meanwhile, a transient expression assay was performed in lily scales by a vacuum penetration method (Sun et al.,2022). Lily scales are usually white to light yellow in colour cultured in the dark and could slightly accumulate anthocyanin transferred to light conditions. When under continuous white light but not under dark conditions, a small amount of anthocyanin was formed after 4 d in the control treated with water (CK) or empty control. However, the presence of LhWRKY44 caused a deeper color than that observed in the control (Figure 9D and E). As expected, further qPCR analysis showed that the addition of LhWRKY44 also activated the transcription of anthocyanin accumulation-related pathway genes, but *UFGT*, *ANS* and *MYBSPLATTER* were highly expressed only under light conditions. Light promoted anthocyanin formation in lily, and this was significantly enhanced in the presence of LhWRKY44.

### LhHY5 activates the transcription of *LhWRKY44*

In our previous study, we found that LhHY5 contributed to the anthocyanin accumulation,and promoted the expression when overexpression(Supplemental Figure S11). To verify that LhHY5 could activates the promoter of LhWRKY44. A dual-luciferase assays was developmented, a construct with the promoter sequences of the LhWRKY44 fused to the LUC reporter vector was combined with the 35S-LhHY5 effector construct for co-infifiltration into tobacco leaves. As shown in Figure. 10A, *LhWRKY44* promoter obviously were activated with the addition of LhHY5, the LUC/REN ratio was relatively higher than that of the control. Moreover, GUS assay showed that GUS staining and the relative expression level of GUS increased when the construct containing the LhHY5 cotransformed into the tepals of lily with the LhWRKY44 promoter (Figure. 10B). To conclude, these data demonstrate that LhHY5 activates the transcription of LhWRKY44.

**Figure 10.**
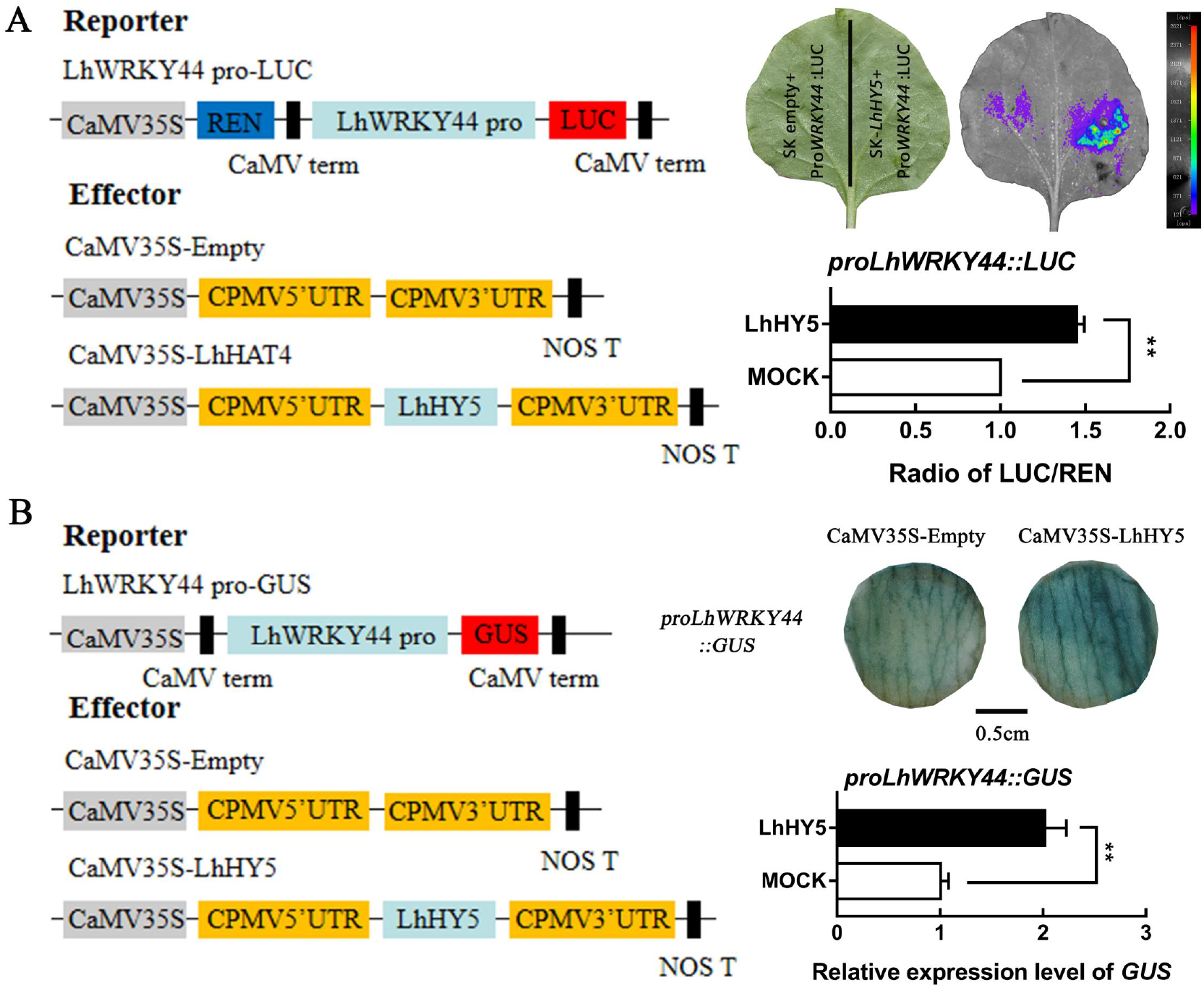
LhHY5 activates the transcription of *LhWRKY44*. A, Dual-luciferase assay of LhHY5 activated *LhWRKY44* transcription. B, GUS activity analysis. GUS staining of lily tepals. Scale bars, 0.5 cm. Mean values ± SDs are shown at least three biological replicates. Asterisks represent T test significance (*P < 0.05,**P < 0.01 and ***, P < 0.001).

## Discussion

### LhWRKY44 positive regulates lily anthocyanin accumulation

Lily is one of high commercial value horticultural crops with divers anthocyanin pigmentation pattern. As a group within the flavonoid family of plant secondary metabolites, Anthocyanins determine the color of flowers, organs and protect plants from reactive oxygen species produced under abiotic stress (Grotewold, 2006). Research over the past few decades has highlighted the critical role of WRKY TFs in plant stress, flowering, development and other processes (Rushton et al., 2010). In addition, WRKY TFs are also recognized as crucial regulators of anthocyanin accumulation, which can function as activators or repressors based on their transcriptional activities.

In *Arabidopsis*, a gene encodes a WRKY44 TF, TTG2 determines trichome formation and seed coat development in the PA pathway (Gonzalez et al., 2016). In *petunia*, the PH3 (WRKY44) TF, which is highly similar to AtTTG2, is activated by the MBW complex,and then activates the transcription of a vacuolar ATPase to hyperacidify the vacuole (Verweij et al., 2016). *VvWRKY26*, a *PhPH3* gene homolog in *Vitis vinifera*, increases the flavonoid content (Amato et al., 2016). Likewise, the kiwifruit WRKY44 regulates the expression of *F3’H* and *F3’5’H* to promote anthocyanin accumulation (Peng et al., 2020). SmWRKY44 could interact with SmMYB1 to jointly promote light-induced anthocyanin accumulation in eggplant (He et al., 2021). However, not all WRKYs have a positive regulatory effect, such as BnWRKY41-1 (WRKY44 homologous gene) from *Brassica napus, which* negatively regulates anthocyanin accumulation in *Arabidopsis* and inhibits the expression of *AtMYB75, AtMYB111* and *AtMYBD*, which are anthocyanin accumulation regulatory genes (Duan et al., 2018). Here, we found a hub gene *LhWRKY44* (c117585_g2) according to coexpression gene network modules constructed using WGCNA during tepal development based on a previous study (Xu et al., 2017). The transcript abundance of the *LhWRKY44* gene showed a significant positive correlation with anthocyanin content (Figure 2; Supplemental Figure S3). Overexpression of *LhWRKY44* promoted anthocyanin accumulation, whereas silencing of *LhWRKY44* decreased anthocyanin accumulation(Figure 3; Supplemental Figure S4 and S5). Concurrently, we showed that LhWRKY44 possesses transcriptional activation activity by the DLR assay (Figure 2C) as described by Han (2016) and Zhang (2021). Thus, LhWRKY44 promotes anthocyanin accumulation by acting as a positive regulator.

### The high conserved and particular roles of the MBW complex

Anthocyanin accumulation in plants is regulated by the MBW protein complex. In lily, a large number of MYB transcription factors have been reported to be involved in anthocyanin biosynthesissuch. Such as, LhMYB12 from Asiatic hybrids lily was identified as the key gene closely related to anthocyanin formation in tepals, filaments and stigmas (Yamagishi et al., 2012; 2014; 2017). LcMYB12, LhSorMYB12, the homologous genes of LhMYB12 play a positive role in regulating the anthocyanin synthesi(s Yamagishi, 2011; 2013; 2020a). LhMYBSPLATTER isolated from Asiatic hybrids lily ‘Latvia’ determines the formation of splashy spots caused by anthocyanin in tepals (Yamagishi et al., 2014a; Yamagishi, 2020a; 2020b). LcMYBspl, a homologous gene of LhMYBSPLATTER, regulates spot pigmentationin *L. cernuum* (Yamagishi et al., 2020a). MYB19long simultaneously regulated the coloring formation of raised spots and brush marks. LhMYB6 is related to spot pigmentation and regulates the coloring of raised spot (Yamagishi et al., 2010). LrMYB15 regulates the coloring of the lower epidermis of the tepals of *L. regale* (Yamagishi, 2016). LhMYB18 controls the formation of spots on the inner surface of tepals in Asiatic hybrids lily (Yamagishi, 2018).

Tango series cultivars are a type of Asiatic hybrid lily and frequently develop red or dark red spots on the interior surfaces of their tepals, and the anthocyanin accumulation within these spots appears on the flowers. Many small splatter spots (speckles) develop in the lower half of their tepals(Yamagishi, 2013b; Yamagishi and Akagi, 2013a). In order to better define the possible regulation anthocyanin biosynthesis functions of MBW complex in Tango series cultivar ‘Tiny Padhye’, we construct a phylogenetic tree after analysing the transcriptome data according to the NR database annotation information of MYBs, bHLHs transcription factor. As shown as Supplementary Figure S6, 8 LhMYBs were divided into subgroup 5 and 6 known MYB regulators in flavonoid pathway. 4 LhbHLHs were clustered into an evolutionary branch on reported LhbHLH1 and LhbHLH2 respectively in the IIIf subgroup containing sequences related to the regulation of anthocyanin biosynthesis according to Feller et al. (2011). qPCR analysis showed that the expression of c51332-g1 (LhMYBSPLATTER) gene was approximately 400 folds higher in the basal parts than in the upper parts among the subgroup 6 R2R3-MYB genes, which is consistent with the statement that LhMYBSPLATTER is mainly responsible for the development of splatter-type spots in the Tango series cultivars of Asiatic hybrid lilies (Yamagishi et al., 2014a;b). And only 115355-g1 (LhbHLH2) showed a expression profiles were consistent with those of the biosynthesis genes among the IIIf bHLH genes. Expression analysis of LhMYBSPLATTER, LhbHLH2 and LhWDR at different development stages and other five Tango series cultivars also revealed that LhMYBSPLATTER, LhbHLH2 and LhWDR may function in anthocyanin accumulation by forming MBW complex.

To verify the above conjecture, we studied the relationship between LhMYBSPLATTER and LhbHLH2. The BiFC and Y2H assays showed that LhMYBSPLATTER interacted with LhbHLH2 to function as an MBW complex member(Supplemental Figure S7D). Similarly, LhWDR was also found to be related to anthocyanin accumulation via interactions with LhbHLH2, consistent with the reports of Dou et al. (2020) (data not shown). These results indicated the highly conserved and particular roles of the MBW complex consisting of LhMYBSPLATTER, LhbHLH2 and LhWDR in anthocyanin accumulation in Asiatic hybrid lilies.

### LhWRKY44 showed a different regulatory network from other WRKY44s

Anthocyanin accumulation is a complex biological process involving multiple enzymes and gene families encoding enzymes such as PAL, CHI, CHS, F3H, DFR, and UFGT. In this study, LhWRKY44 was characterized as a positive regulator of anthocyanin accumulation in lily (Figure 3). Since the lily genome has not been sequenced, we identified the promoter sequences of structural genes and regulatory genes using the narrow homologous cloning and chromosome walking techniques with specific primers designed according to Lai (2012) and Cao (2021a;2021b). Y1H assay and EMSAs demonstrated that LhWRKY44 promotes anthocyanin accumulation by directly binding to the *PAL* and *F3H* promoters. The dual-luciferase assays also showed that LhWRKY44 could activate the expression of *LhCHS, LhDFR, LhUFGT* promoters, LhWRKY44 may function on these genes indirectly.

Research suggests the WRKY-MBW module play an essential role in regulating flavonoid accumulation. In *Petunia*, the regulatory network of the PH3(WRKY44) TF only shown to join the PH4-AN1-AN11 (MBW) complex by interacting with the WD40 protein AN11, and activated the expression of *PH5* to regulate vacuolar acidification for anthocyanin storage(Verweij et al., 2016). Similarly, in *Arabidopsis*, the TTG2 interacts with the WD40 protein TTG1 of the MBW complex to regulated TT12 and TT13/AHA10 genes involved in transporting pigment precursors into the vacuole of inner testa cells (Gonzalez et al., 2016). Phylogenetic analysis showed that LhWRKY44 had a close evolutionary relationship with PH3 and TTG2, which means that they may have a similar regulatory mechanism (Figure 1A). Therefore, we investigated whether the same mechanism also exists in lily. First, we isolated PH5 (a homolog of AHA10) based on transcriptome data and analyzed its expression in *LhWRKY44-overexpressing and LhWRKY44-silenced* tepals, and this did not trigger a change in *pH5* expression (Supplemental Figure S10A). Meanwhile, there was no significant difference in pH5 expression between the upper and basal tepals and no obvious correlation with anthocyanin accumulation in lily as shown in Supplemental Figure (S10B). Then, we explored the relationship between the LhWRKY44 and LhWDR proteins. Simultaneously, the results showed that the LhWRKY44 and LhWDR proteins did not interact (Figure 6A and B). LhWRKY44 has a different and unique regulatory mechanism with those of PH3 and TTG2(Gonzalez et al., 2016; Verweij et al., 2016). Interestingly, here, the novel dual roles of the WRKY-MBW regulatory module may play an essential role in regulating anthocyanin accumulation in lily, in which LhWRKY44 not only binds to *LhMYBSPLATTER* but also interact with LhMYBSPLATTER and enhance its transcriptional activation of its target genes and the formation of MBW complex (Figure 7A and B). These results suggest that anthocyanin production is a finely tuned biological process involving different TFs.

It is worth mentioning that our results proved that LhWRKY44 could directly bind to *LhGST* (Figure 5), an intracellular transport protein involved in anthocyanin accumulation, which is unique and distinctive to other reported WRKY44 TF regulation modules of anthocyanin accumulation in plants. Multipathway regulation of LhWRKY44 is associated with anthocyanin accumulation, transport and transcriptional level, which is the first reported mechanism of the role of WRKY TFs in the transcription of transport-related genes in lily up to now.

### LhWRKY44 is central TF of the light-induced anthocyanin biosynthesis cascade

Anthocyanins accumulation in plant tissues and flowers is jointly controlled by internal factors and external environmental factors. Light, temperature, hormones, sugar, etc., affect the accumulation of anthocyanins (Jiang et al., 2014; An et al., 2019). Much progress has been made toward light promotes anthocyanin accumulation in lily. (Nakatsuka et al., 2009;Yamagishi et al., 2016). The WRKY TF superfamily participate in numerous physiological processes, including flowering, development, senescence and stress responses). WRKY TFs are also identified to participate in light signaling pathways. In *Arabidopsis*, AtWRKY40 and AtWRKY63 activate the transcription of light signaling pathway genes (Aken et al., 2013). In apple, light induces *MdWRKY1* expression, and promoted anthocyanin accumulation(Ma et al., 2021).

Here, we proved LhWRKY44 is a component of light signaling. The promoter analysis of *LhWRKY44* displayed a large number of light-responsive elements, such as the GT1-motif, Sp1, TCT-motif, AE-box, LAMP-element, and Gap-box except core elements, TATA box, CAAT box, and G-box. The expression of *LhWRKY44* was induced when lily were exposed to light (Figreu. 9). Notably, HY5 regulate other TFs and affect photomorphogenesis and light-induced anthocyanin accumulation at the transcriptional and post-translational levels in *Arabidopsis* (Datta et al., 2008). and HY5 positively mediates photomorphogenesis by regulating a light-responsive element G-box (Bai et al., 2019). Thus, it would be interesting to investigate whether LhWRKY44 targeted by HY5 gene associated with the light response pathway to coregulate anthocyanin accumulation-related genes. A *LhHY5* could be induced by light and showed positive correlation with athocyanin content. And highly significant expression of LhWRKY44 was detected in tepal overexpressing LhHY5 (Supplemental Figure S11), suggesting that LhHY5 might regulate LhWRKY44 expression. Dual-luciferase assay and GUS report assay revealed that LhHY5 induce the expression of *LhWRKY44* (Figure 10). Although further detailed and direct interaction in vivo experiments are needed, these results generate a central-LhWRKY44 cascade (LhHY5-LhWRKY44-anthocyanin pathway gense) involved in light-mediated anthocyanin accumulation in Asiatic hybrid lilies.

## Conclusion

In conclusion, we revealed the molecular mechanism by LhWRKY44 to regulate anthocyanin accumulation. LhWRKY44 activated the transcription of *LhPAL*, *LhF3H* and a intracellular anthocyanin transport protein LhGST. Additionally, the novel dual role of WRKY-MBW of which LhWRKY44 interacts with LhMYBSPLATTER, strongly promoted with the formation of MBW complex, and indirectly enhance the expression of *DFR*, *UFGT* and *GST*, which are targeted by LhMYBSPLATTER, thus promoting anthocyanin accumulation (Figure 11).

**Figure 11.**
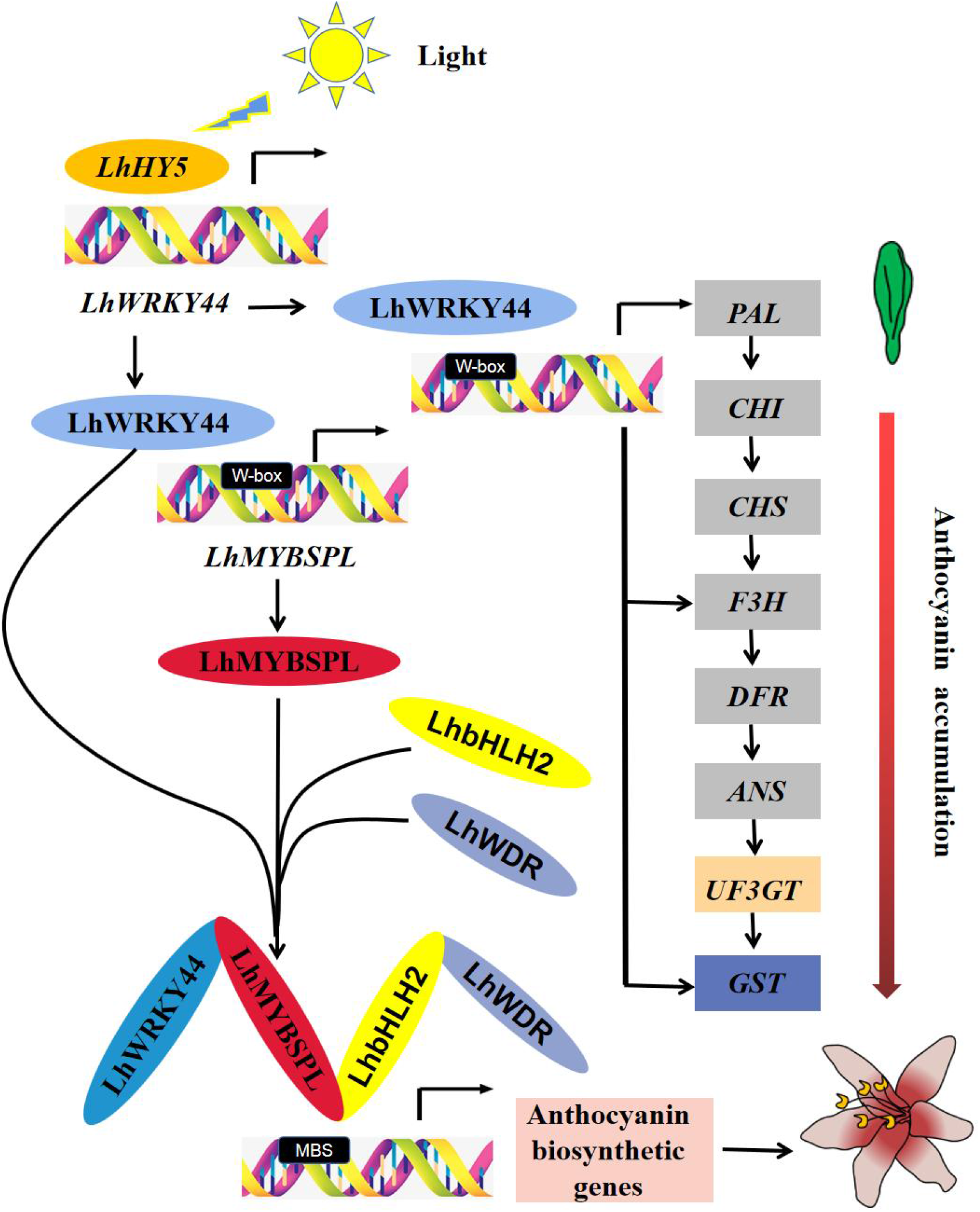
A proposed working model of LhWRKY44 regulatory network in anthocyanin accumulation in lily.

## Materials and methods

### Plant materials, growth conditionsand treatments

Lily cultivars were grown in a greenhouse located in the Institute of Vegetables and Flowers, Chinese Academy of Agricultural Sciences (Beijing, China). Flower samples were collected at five stages defined according to Xu et al. (2017) with minor modifications. Tobacco (*Nicotiana tabacum and Nicotiana benthamiana*) plants were grown in nutritive soil with 70% relative humidity in a programmable incubator with a 16 h photoperiod at 22°C. Tobacco was used in the present study for the transformation experiments.

The method used to determine the effect of light on anthocyanins in lily was followed Nakatsuka et al. (2009). ‘Tiny Padhye’ with 1-2-cm-long flower buds were grown in dark and high-light (60000 lux) conditions for 12 d in an incubator at 23°C. All these samples were immediately chopped, frozen and stored at −80°C.

### Total RNA isolation, cDNA accumulation and RT-qPCR analysis

Total RNA was extracted by using an RNA Prep Pure Plant Kit (polysaccharide and polyphenolic-rich) (TIANGEN, Beijing, China) following the manufacturer’s instructions. TransScript^®^ II One-Step gDNA Removal and cDNA Accumulation SuperMix (TransGen Biotech, Beijing, China) was used for cDNA synthesized. Hieff^®^ qPCR SYBR^®^ Green Master Mix (YEASEN, Shanghai, China) was used for RT-qPCR. For each sample, quantifications were performed. The relative expression levels was calculated with 2^-ΔΔCt^ method. The primers used for qPCR are listed in Supplemental Table S2.

### Anthocyanin extraction and measurement

The total anthocyanin extraction and measurement was determined followed by Cao et al. (2021). There were at least three biological replicates per sample. The total anthocyanin content was calculated according to the formula: OD = (A530 - A620) - 0.1 × (A650 - A620).

### RACE assay and bioinformatics analysis

LhWRKY44 full-length was amplified by the GeneRacer kit (Invitrogen, Carlsbad, CA, USA). The primer used are shown in Supplemental Table S3. Multiple sequence alignments were conducted by DNAMAN with default parameters. Using MEGA software version 6 constructed the phylogenetic trees with the neighbor-joining (NJ) algorithm method and 1,000 bootstrap replicates.

### Transcriptional activity analysis

Vector construction and transactivation activity assay of LhWRKY44 in tobacco leaves were carried out as described by Han et al. (2016) and Zhang et al. (2021), LhWRKY44 was inserted to pBD vector to construct the effector plasmid. The reporter harbored a LUC reporter gene, with a REN reporter driving by a minimal 35S promoter as an internal control. The effector and reporter vectors were co-infiltration into *N. benthamiana*. After 3 d, measured the ratios of LUC to REN by the Dual Luciferase Reporter Gene Assay Kit (Yeason, Shanghai, China). The activity was analyzed with at least three biological replications. LUC signals living images were captured by a low-light cooled CCD imaging apparatus.

### Subcellular localization

The CDS of LhWRKY44 without the stop codon was inserted into a pCAMBIA2300-eGFP vector to generate a fusion protein and then transformed into *A. tumefaciens* strain GV3101. GFP construction primers are shown in Supplemental Table S4. The bacterial solution harbored LhWRKY44-eGFP and eGFP empty plasmid(control) were infiltrated into four-week-old *N. benthamiana* tobacco leaves. The GFP signals in the leaf epidermal cells were imaged after 3 d infiltration in four-week-old tobacco leaves.

### Transient overexpression and virus-induced gene silencing of *LhWRKY44* in lily tepals

To confirm the function of LhWRKY44, we transiently overexpressed LhWRKY44 in the lily cultivar ‘Robina’ and silenced it in ‘Tiny Padhye’ basal tepals according to Fatihah et al. (2019) and Yin et al. (2021). Briefly, the outer tepals of lily flower buds of ‘Robina’ (approximately 6-7 cm), which were green and just beginning to open, were infiltrated. For the generation of pTRV2-LhWRKY44, a specific fragments of ~240 bp of LhWRKY44 were PCR amplified and infused into the pTRV2 vector (Supplemental Table S4). Agrobacterium EHA105 cultures containing combinations of pTRV1, pTRV2, or its derivatives were mixed in a 1:1:ratio before infiltration. Then, they were introduced into the colored buds of the tepals close to stage 3 (whose lengths were approximately 4-5 cm). There were 10 plants per replicate in each treatment and at least three biological replicates. Taking the digital photographs after 1 week.

### Dual-luciferase reporter (DLR) assay

Dual-luciferase reporter (DLR) assays were performed as previously described (He et al., 2022), where a pGreenII 0029 62-SK vector was fused to the CDS of LhWRKY44 as the effector and with a pGreenII 0800-LUC vector carrying the promoter sequences as reporters. DLR assay primers sequences are shown in Supplemental Table S4. The pGreenII 0029 62-SK empty vector was used as control. Subsequently, the GV3101 bacterial solution containing the reporter and effector constructs were transiently introduced into tobacco leaves. The luciferase signals were imaged and measured as mentioned above.

### Yeast one-hybrid (Y1H) assay

Y1H assays were performed using the Matchmaker Gold Yeast One-Hybrid Library Screening System (Clontech, USA). The sequence of LhWRKY44 was inserted into the vector pGADT7 as prey. The promoters were inserted into the pAbAi vector as bait, then transformated them into the Y1H Gold yeast strain, the transformants were spread on SD/-Ura/AbA (aureobasidin A) culture medium for autoactivation tests. Primers used in Y1H assays are listed in Supplemental Table S4. Subsequently, the interaction physical interaction was determined according to colony growth on SD/-Leu medium supplemented with corresponding AbA concentrations, allowing the colonies to grow for 3-5 d at 28°C.

### Electrophoretic mobility shift assay (EMSA)

To obtain soluble protein, the coding sequence of LhWRKY44 was cloned into the vector pET-32a (with 6× His-tag) to produce recombinant LhWRKY44 protein with a polyhistidine (His) tag in *Escherichia coli* strain BL21 (DE3). Ni-NTA SefinoseTM Resin (Sangon, Shanghai) was subsequently used to purify the proteins. EMSA was performed using the LightShift™ Chemiluminescent EMSA Kit (Thermo Fisher Scientific, USA) according to the manufacturer’s protocol. The specific primer sequences used in the EMSAs are listed in Supplemental Table S4.

### Bimolecular fluorescence complementation assay (BiFC)

The LhWRKY44 and LhMYBSPLATTER CDSs lacking stop codons were inserted into the P2YC and P2YN vectors, respectively. The primers are presented in Supplemental Table S4. Then the GV3101 bacterial solution containing the resultant vectors with the nuclear marker co-infiltrated into *N. benthamiana* leaves as previously described (Ni et al., 2021) were introduced into *A. tumefaciens* strain GV3101 and and visualized by high-resolution confocal laser microscopy after 3 d of transformation.

### Yeast two-hybrid (Y2H) assay

The sequence of LhWRKY44 was amplied into pGADT7 as prey vector. The sequences of LhMYBSPLATTER, LhbHLH2 and LhWDR were amplied into pGBKT7 as bait vectors. Y2H assay was performed by the manufacturer’s protocol of the Matchmaker GAL4 Two-hybrid System (Clontech, USA). The prey and bait were cotransformed into *Saccharomyces cerevisiae* strain AH109. Protein-protein interactions were detected on DDO (SD medium lacking Leu and Trp) and QDO/X (SD medium lacking Leu, Trp, His and Ade, with X-α-gal) media.

### Firefly luciferase complementation imaging (FLC) assay

The vector information and protocol for the FLC assay were based on the studies of Shi et al. (2021). Additionally, LhWRKY44 was incorporated into the pGreenII 0029 62-SK vector for subsequent overexpression. The recombinant plasmids were transformed into GV3101, after which different combinations comprising equal volumes of each vector were used to infiltrate tobacco leaves as mentioned above. Then the LUC signal was imaged by a CCD imaging camera.

### GUS staining assays

The promoter of LhWRKY44 was cloned using a Genome Walker Kit (Clontech) with nested PCR from genomic DNA of ‘Tiny Padhye’, with the primers listed in Supplementary Table S2. Promoter sequence was analyzed by PLANTCARE software. Then LhWRKY44 promoter was introduced into the reporter pBI121 vector harbored the GUS report gene. The CDS of LhHY5 inserted into pGreen0600-SK, creating the effector vector, then transferred into A. tumefaciens strain EHA105. Tepals at stage 3 were used for infiltration. Three days after infiltration, the tepals were treated with GUS staining solution (Solarbio, China), washed and cleared with 70% ethanol for more than 24 h before image capture. Relative GUS expression level were analysed by RT-qPCR.

### Accession numbers

Gene accession numbers used in this study: *LhWRKY44* (OP380462), *LhPAL(* AB699158), *LhCHI*(KJ784468), *LhCHS*(AB715424), *LhF3’H(AB699162), LhF3H*(AB699169), *LhDFR*(AB699165), *LhANS* (AB699166),*LhUFGT*(KX781249), *LhGST*(MK426728), *LhMYBSPLATTER*(MW719045), *LhbHLH*2(AB222076), *LhWDR*(MW509631), *LhHY5*(OQ121997), *LhPH5* (OQ121998).

## Supplemental data

The following materials are available in the online version of this article.

**Supplemental Figure S1.** Multiple sequence alignment of the LhWRKY44 transcription factor.

**Supplemental Figure S2.** Phylogenetic tree analysis of LhWRKY44 and other WRKY proteins from *A. thaliana*.

**Supplemental Figure S3.** Different tissues and tepal development stages of the Asiatic hybrid lily cultivar ‘Tiny Padhye’.

**Supplemental Figure S4.** Diagrams of sampling positions in the infiltrated sites in lily tepals with overexpression and silencing.

**Supplemental Figure S5.** Promoters sequence analysis.

**Supplemental Figure S6.** Phylogenetic trees of LhMYBs and LhbHLHs in ‘Tiny Padhye’.

**Supplemental Figure S7.** MBW complex of LhMYBSPLATTER, LhbHLH2 and LhWDR highly conserved and particular roles in ‘Tiny Padhye’.

**Supplemental Figure S8.** Direct binding of LhWRKY44 to the promoters of *LhMYBSPLATTER*.

**Supplemental Figure S9.** The expression level of anthocyanin pathway genes after high light treatment.

**Supplemental Figure S10.** Expression analysis of *LhPH5*.

**Supplemental Figure S11.** Expression analysis of *LhHY5*.

## Acknowledgments

We thank Prof. Nan Ma, Prof. Xia Cui, Prof. Allen A C, Prof. Yuanwen Teng, Prof. Ji Tian for assistance with providing experimental plasmids and guidances.

## Reference

Amato A, Cavallini E, Zenoni S, Finezzo L, Begheldo M, Ruperti B, Tornielli GB (2016) A grapevine TTG2-Like WRKY transcription factor is involved in regulating vacuolar transport and flavonoid biosynthesis. Front. Plant Sci. 7: 1979.

An JP, Zhang XW, You CX, Bi SQ, Wang XF, Hao YJ (2019) MdWRKY40 promotes wounding-induced anthocyanin biosynthesis in association with MdMYB1 and undergoes MdBT2-mediated degradation. New Phytol. 224: 380–395.

Bai S, Tao R, Tang Y, Yin L, Ma Y, Ni J, Yan X, Yang Q, Wu Z, Zeng Y, et al. (2019) BBX16, a B-box protein, positively regulates light-induced anthocyanin accumulation by activating MYB10 in red pear. Plant Biotechnol. J. 17: 1985–1997.

Cao Y, Bi M, Yang P, Song M, He G, Wang J, Yang Y, Xu L, Ming J (2021) Construction of yeast one-hybrid library and screening of transcription factors regulating LhMYBSPLATTER expression in Asiatic hybrid lilies (*Lilium* spp.). BMC Plant Biol. 21: 563.

Cao Y, Xu L, Xu H, Yang P, He G, Tang Y, Qi X, Song M, Ming J (2021) LhGST is an anthocyanin-related glutathione S-transferase gene in Asiatic hybrid lilies (*Lilium* spp.). Plant Cell Rep. 40: 85–95.

Datta S, Johansson H, Hettiarachchi C, Irigoyen ML, Desai M, Rubio V, Holm M (2008) LZF1/SALT TOLERANCE HOMOLOG3, an *Arabidopsis* B-box protein involved in light-dependent development and gene expression, undergoes COP1-mediated ubiquitination. Plant Cell. 20: 2324–2338.

Dou X, Bai J, Wang H, Kong Y, Lang L, Bao F, Shang H (2020) Cloning and characterization of a tryptophan–aspartic acid repeat gene associated with the regulation of anthocyanin biosynthesis in Oriental hybrid lily. J. Amer.Soc.Hort.Sci. 145: 131–140.

Duan S, Wang J, Gao C, Jin C, Li D, Peng D, Du G, Li Y, Chen M (2018) Functional characterization of a heterologously expressed *Brassica napus* WRKY41-1 transcription factor in regulating anthocyanin biosynthesis in *Arabidopsis thaliana*. Plant Sci. 268: 47–53.

Fatihah HNN, Moñino López D, van Arkel G, Schaart JG, Visser RGF, Krens FA (2019) The ROSEA1 and DELILA transcription factors control anthocyanin biosynthesis in *Nicotiana benthamiana* and *Lilium* flowers. Sci. Horti. 243: 327–337.

Feller A, Machemer K, Braun EL, Grotewold E (2011) Evolutionary and comparative analysis of MYB and bHLH plant transcription factors. Plant J. 66: 94–116.

Gonzalez A, Brown M, Hatlestad G, Akhavan N, Smith T, Hembd A, Moore J, Montes D, Mosley T, Resendez J, et al. (2016) TTG2 controls the developmental regulation of seed coat tannins in *Arabidopsis* by regulating vacuolar transport steps in the proanthocyanidin pathway. Dev Biol. 419: 54–63.

Grotewold E (2006) The genetics and biochemistry of floral pigments. Annu Rev Plant Biol. 57: 761–780.

Han YC, Kuang JF, Chen JY, Liu XC, Xiao YY, Fu CC, Wang JN, Wu KQ, Lu WJ (2016) Banana transcription factor MaERF11 recruits histone deacetylase MaHDA1 and represses the expression of *MaACO1* and expansins during fruit ripening. Plant Physiol. 171: 1070–1084.

He G, Cao Y, Wang J, Song M, Bi M, Tang Y, Xu L, Ming J, Yang P (2022) WUSCHEL-related homeobox genes cooperate with cytokinin to promote bulbil formation in *Lilium lancifolium*. Plant Physiol. 190: 387–402.

He Y, Wang Z, Ge H, Liu Y, Chen H (2021) Weighted gene co-expression network analysis identifies genes related to anthocyanin biosynthesis and functional verification of hub gene SmWRKY44. Plant Sci. 309: 110935.

Jiang Y, Liang G, Yang S, Yu D (2014) *Arabidopsis* WRKY57 functions as a node of convergence for jasmonic acid- and auxin-mediated signaling in jasmonic acid-induced leaf senescence. Plant Cell. 26: 230–245.

Lai YS, Shimoyamada Y, Nakayama M, Yamagishi M (2012) Pigment accumulation and transcription of *LhMYB12* and anthocyanin biosynthesis genes during flower development in the Asiatic hybrid lily (*Lilium* spp.). Plant Sci. 193-194: 136–147.

Li C, Wu J, Hu KD, Wei SW, Sun HY, Hu LY, Han Z, Yao GF, Zhang H (2020a) PyWRKY26 and PybHLH3 cotargeted the PyMYB114 promoter to regulate anthocyanin biosynthesis and transport in red-skinned pears. Hortic Res. 7: 37.

Liu W, Wang Y, Yu L, Jiang H, Guo Z, Xu H, Jiang S, Fang H, Zhang J, Su M, et al. (2019) MdWRKY11 participates in anthocyanin accumulation in red-fleshed apples by affecting MYB transcription factors and the photoresponse factor MdHY5. J Agric Food Chem. 67: 8783–8793.

Ma H, Yang T, Li Y, Zhang J, Wu T, Song T, Yao Y, Tian J (2021) The long noncoding RNA MdLNC499 bridges MdWRKY1 and MdERF109 function to regulate early-stage light-induced anthocyanin accumulation in apple fruit. Plant Cell. 33: 3309–3330.

Nakatsuka A, Izumi Y, Yamagishi M (2003) Spatial and temporal expression of chalcone synthase and dihydroflavonol 4-reductase genes in the Asiatic hybrid lily. Plant Sci. 165: 759–767.

Nakatsuka A, Yamagishi M, Nakano M, Tasaki K, Kobayashi N (2009) Light-induced expression of basic helix-loop-helix genes involved in anthocyanin biosynthesis in flowers and leaves of Asiatic hybrid lily. Sci. Hortic. 121: 84–91.

Nakatsuka T, Haruta KS, Pitaksutheepong C, Abe Y, Kakizaki Y, Yamamoto K, Shimada N, Yamamura S, Nishihara M (2008) Identification and characterization of R2R3-MYB and bHLH transcription factors regulating anthocyanin biosynthesis in gentian flowers. Plant Cell Physiol. 49: 1818–1829.

Ni J, Bai S, Zhao Y, Qian M, Tao R, Yin L, Gao L, Teng Y (2019) Ethylene response factors Pp4ERF24 and Pp12ERF96 regulate blue light-induced anthocyanin biosynthesis in ‘Red Zaosu’ pear fruits by interacting with MYB114. Plant Mol Biol. 99: 67–78.

Ni J, Premathilake AT, Gao Y, Yu W, Tao R, Teng Y, Bai S (2021) Ethylene-activated *PpERF105* induces the expression of the repressor-type R2R3-MYB gene *PpMYB140* to inhibit anthocyanin biosynthesis in red pear fruit. Plant J. 105: 167–181.

Rushton PJ, Somssich IE, Ringler P, Shen QJ (2010) WRKY transcription factors. Trends in Plant Sci. 15: 247–258.

Sakai M, Yamagishi M, Matsuyama K (2019) Repression of anthocyanin biosynthesis by R3-MYB transcription factors in lily (Lilium spp.). Plant Cell Rep. 38: 609–622.

Shi Y, Pang X, Liu W, Wang R, Su D, Gao Y, Wu M, Deng W, Liu Y, Li Z (2021) SlZHD17 is involved in the control of chlorophyll and carotenoid metabolism in tomato fruit. Hortic Res. 8: 259.

Sun L, Song Sl, Yang Y, Sun HM (2022) Melatonin regulates lily bulblet development through the LoBPM3-LoRAV module. Ornamental Plant Res. 2: 1–11.

Sun Y, Li H, Huang JR (2012) Arabidopsis TT19 functions as a carrier to transport anthocyanin from the cytosol to tonoplasts. Mol Plant. 5: 387–400.

Suzuki K, Suzuki T, Nakatsuka T, Dohra H, Yamagishi M, Matsuyama K, Matsuura H (2016) RNA-seq-based evaluation of bicolor tepal pigmentation in Asiatic hybrid lilies (*Lilium* spp.). BMC Genomics 17: 611.

Van Aken O, Zhang B, Law S, Narsai R, Whelan J (2013) AtWRKY40 and AtWRKY63 modulate the expression of stress-responsive nuclear genes encoding mitochondrial and chloroplast proteins. Plant Physiology. 162: 254–71.

Verweij W, Spelt CE, Bliek M, de Vries M, Wit N, Faraco M, Koes R, Quattrocchio FM (2016) Functionally similar WRKY proteins regulate vacuolar acidification in *Petunia* and hair development in *Arabidopsis*. Plant Cell. 28: 786–803.

Winkel-Shirley, B (2001) Flavonoid biosynthesis. A colorful model for genetics, biochemistry, cell biology, and biotechnology. Plant Physiol. 126: 485–493.

Xiang L, Liu X, Li H, Yin X, Grierson D, Li F, Chen K (2019) CmMYB#7, an R3 MYB transcription factor acts as a negative regulator of anthocyanin biosynthesis in *Chrysanthemum*. J. Exp. Bot. 70: 3111–3123.

Xu H, Yang P, Cao Y, Tang Y, He G, Xu L, Ming J (2020) Cloning and functional characterization of a flavonoid transport-related MATE gene in Asiatic hybrid lilies (*Lilium* spp.). Genes. 11: 418.

Xu L, Yang P, Feng Y, Xu H, Cao Y, Tang Y, Yuan S, Liu X, Ming J (2017) Spatiotemporal transcriptome analysis provides insights into bicolor tepal development in *Lilium* “Tiny Padhye”. Front Plant Sci. 8: 398.

Xu W, Grain D, Bobet S, Le Gourrierec J, Thevenin J, Kelemen Z, Lepiniec L, Dubos C (2014) Complexity and robustness of the flavonoid transcriptional regulatory network revealed by comprehensive analyses of MYB-bHLH-WDR complexes and their targets in *Arabidopsis* seed. New Phytol. 202: 132–144.

Yamagishi M (2013b) How genes paint lily flowers: Regulation of colouration and pigmentation patterning. Sci. Hortic. 163: 27–36.

Yamagishi M (2016) A novel R2R3-MYB transcription factor regulates light-mediated floral and vegetative anthocyanin pigmentation patterns in *Lilium* regale. Mol. Breeding. 36: 3.

Yamagishi M (2018) Involvement of a LhMYB18 transcription factor in large anthocyanin spot formation on the flower tepals of the Asiatic hybrid lily (*Lilium*spp.) cultivar “Grand Cru”. Mol. Breeding. 38: 60.

Yamagishi M (2020a) Isolation and identification of MYB transcription factors (MYB19Long and MYB19Short) involved in raised spot anthocyanin pigmentation in lilies (*Lilium* spp.). J Plant Physiol. 250: 153164.

Yamagishi M (2020b) White with partially pink flower color in *Lilium cernuum* var. album is caused by transcriptional regulation of anthocyanin biosynthesis genes. Sci. Horti. 260: 108880

Yamagishi M (2020c) MYB19LONG is involved in brushmark pattern development in Asiatic hybrid lily (*Lilium* spp.) flowers. Sci.Horti. 272: 109570.

Yamagishi M, Akagi K (2013b) Morphology and heredity of tepal spots in Asiatic and oriental hybrid lilies (*Lilium* spp.). Euphytica. 194: 325–334.

Yamagishi M, Ihara H, Arakawa K, Toda S, Suzuki K (2014a) The origin of the LhMYB12 gene, which regulates anthocyanin pigmentation of tepals, in Oriental and Asiatic hybrid lilies (*Lilium* spp.). Sci.Horti. 174: 119–125.

Yamagishi M, Nakatsuka T (2017) LhMYB12, Regulating tepal anthocyanin pigmentation in Asiatic hybrid lilies, is derived from *Lilium dauricum* and *L. bulbiferum*. Horti. J. 86: 528–533.

Yamagishi M, Shimoyamada Y, Nakatsuka T, Masuda K (2010) Two R2R3-MYB genes, homologs of Petunia AN2, regulate anthocyanin biosyntheses in flower tepals tepal spots and leaves of asiatic hybrid lily. Plant Cell Physiol. 51: 463–474.

Yamagishi M, Toda S, Tasaki K (2014b) The novel allele of the LhMYB12 gene is involved in splatter-type spot formation on the flower tepals of Asiatic hybrid lilies (*Lilium* spp.). New Phytol. 201: 1009–1020.

Yamagishi M, Yoshida Y, Nakayama M (2011) The transcription factor LhMYB12 determines anthocyanin pigmentation in the tepals of Asiatic hybrid lilies (*Lilium* spp.) and regulates pigment quantity. Mol. Breeding 30: 913–925.

Yin X, Zhang Y, Zhang L, Wang B, Zhao Y, Irfan M, Chen L, Feng Y (2021) Regulation of MYB transcription factors of anthocyanin synthesis in lily flowers. Front Plant Sci. 12: 761668.

Zentgraf U, Laun T, Miao Y (2010) The complex regulation of WRKY53 during leaf senescence of *Arabidopsis thaliana*. Eur J Cell Biol. 89: 133–137.

Zhang Y, Wu Z, Feng M, Chen J, Qin M, Wang W, Bao Y, Xu Q, Ye Y, Ma C, et al. (2021) The circadian-controlled PIF8-BBX28 module regulates petal senescence in rose flowers by governing mitochondrial ROS homeostasis at night. Plant Cell. 33: 2716–2735.

Zhang Z, Shi Y, Ma Y, Yang X, Yin X, Zhang Y, Xiao Y, Liu W, Li Y, Li S, et al. (2020) The strawberry transcription factor FaRAV1 positively regulates anthocyanin accumulation by activation of FaMYB10 and anthocyanin pathway genes. Plant Biotechnol.J. 18: 2267–2279.

